# Edge computer vision produces microarthropod-based high-throughput biodiversity metrics

**DOI:** 10.1101/2025.09.09.675044

**Authors:** Miklós Dombos, Tamara R. Hartke, Zsolt Tóth, Gergő B. Békési, Christoph Scherber, Dénes Schmera, Bernát Zawiasa, László Sipőcz, Veronika Gergócs-Winkler, Norbert Flórián, Vera C. Prenzel, Justine Lejoly, Max Newbert, Jenny Bussell, Luca Serrati, Andrea Veres, András Juhász, Katalin Juhos, Riina Kaasik, Vasileios Vasileiadis

**Affiliations:** HUN-REN Centre for Agricultural Research, Institute for Soil Sciences, Fehérvári Street 132-144. 1116 Budapest, Hungary; Department of Automation and Applied Informatics, Faculty of Electrical Engineering and Informatics, Budapest University of Technology and Economics, Műegyetem rkp. 3., H-1111 Budapest, Hungary; Edaphone Ltd., Berényi Zsigmond street 6, 2500 Esztergom, Hungary; Syngenta (United Kingdom): Fulbourn, England, GB; The Allerton Project, Game and Wildlife Conservation Trust, Game and Wildlife Conservation Trust, Burgate Manor, Fordingbridge, SP6 1EF; Syngenta Italia S.p.A., Viale Fulvio Testi 280/6; Leibniz Institute for the Analysis of Biodiversity Change, Museum Koenig, Adenauerallee 127, 53113 Bonn, Germany; Netherlands Institute of Ecology (NIOO-KNAW), Department of Terrestrial Ecology; HUN-REN Balaton Limnological Research Institute, Klebelsberg K. u. 3, H-8237 Tihany, Hungary; Department of Integrated Plant Protection, Hungarian University of Agriculture and Life Sciences; Pater K. str. 1., H-2100 Gödöllő, Hungary; Department of Agro-Environmental Studies, Hungarian University of Agriculture and Life Sciences; Villányi str. 29-43, H-1118 Budapest, Hungary; Estonian University of Life Sciences, F. R. Kreutzwaldi 1, 50410, Tartu, Estonia; Syngenta Crop Protection AG; Syngenta Crop Protection AG, Rosentalstrasse 67, Basel 4058, Switzerland; Bonn Institute for Organismic Biology, University of Bonn, Germany

## Abstract

Soil fauna is crucial for carbon cycling, controlling organic matter decomposition and contributing to ecosystem services^1–7^. Soil microarthropods are top-down regulators in the decomposer food web and serve as fundamental indicators of soil health. Yet, routine field-level monitoring remains restricted to resource-intensive research, leaving a critical gap in arable land management and policy. To address this, we developed Edapholog^®^ extractor, a fully automated laboratory device that detects and identifies live soil microarthropods via real-time, AI-based image analysis, enabling taxonomic identification. Here, we demonstrate its accuracy across 319 arable fields spanning ten European countries. We show that computer vision can support ecological interpretation in arable systems and long-term studies on conservation tillage and cover cropping, offering a scalable tool for integrating soil biodiversity metrics into regenerative agriculture and carbon farming. We validated the system against classical taxonomy and found that across ∼35,000 microarthropod individuals, the device achieved an overall accuracy of 86%, sensitivity of 75%, and specificity of 99% compared to manual identifications. Community composition analyses revealed high similarities (83%), with minimal richness differences (7%) and low species replacement (13%) across countries, indicating that the AI does not introduce taxonomic bias. When applied in a long-term field experiment, the system detected significant taxon-specific responses to conservation tillage, with effect sizes ranging from 0.5 to 4. Total abundance, richness, and a soil biological health index were 39%, 47%, and 150% higher, respectively, under conservation tillage compared to conventional ploughing. These effects were statistically consistent between the automated and classical methods. However, while manual microscopy required several hours per sample, the AI-based system delivered immediate results without the need for taxonomic expertise. Edapholog^®^ extractor offers exciting opportunities for rapid, scalable soil biodiversity monitoring for future sustainable land management.

## 1. Introduction

Soil, a living medium, encompasses ecological processes providing ecosystem services, most of which are intricately linked to soil organisms^8,9^. The crucial role of soil-dwelling invertebrates, including soil mesofauna (microarthropods) in organic matter (humus) development underscores the significance of this research area^2–4^. In soil ecology, the current focus is on developing and establishing effect-based soil biodiversity monitoring systems able to track soil biological responses to environmental loads and restoration actions on both local and global scale^10–14^. Regenerative agriculture is gaining unprecedented momentum, driven by carbon programs aimed at increasing soil carbon stocks through enhanced humus formation. Practices such as reduced tillage, retention of crop residues, mulching, organic fertilization, and cover cropping not only facilitate carbon sequestration but also invigorate soil biological activity and enrich biodiversity - essential for decomposition and stable humus formation^15–19^. Reflecting this trend, international initiatives and policy frameworks, including the European Union’s Soil Monitoring Law^20^, increasingly emphasize comprehensive monitoring of soil biodiversity. Programs such as the Soil Biodiversity Observation Network (SOILBON) exemplify global efforts to systematically track soil biota^12,21^; however, current recommended methodologies are often constrained by labour intensity and taxonomic limitations, underscoring the urgent need for innovative, scalable approaches to accurately assess regenerative practices and inform sustainable land management policies^12,13,22^. Emerging monitoring methods such as environmental DNA and remote sensing are recommended for assessing soil and ecosystem health^23–25^. Concurrently, soil biodiversity is gaining recognition in environmental finance and corporate accountability: it is increasingly discussed as a potential basis for biodiversity and carbon credits^26,27^. The European Union’s Corporate Sustainability Reporting Directive (CSRD) mandates biodiversity impact disclosures under the ESRS E4: Biodiversity and Ecosystems standard, where soil biodiversity is explicitly identified as a key indicator.

Over the past decade, camera trap technologies have been widely adopted to detect flagship species in conservation biology and to monitor insect pests in plant protection, with recent advances largely focused on image-based analysis^28^. In the case of microarthropods, in addition to previously used infrared gate solutions^29–32^, AI-driven image processing approaches have also emerged, primarily targeting preserved and captured specimen^33–35^. These solutions typically integrated object detection with classification using either the then-current YOLO variants or region-based two-stage methods such as Faster R-CNN, or applied watershed-based segmentation on high-resolution images followed by CNN classification with DenseNet, ResNet, or MobileNet models. These systems have so far been applied primarily in case-study contexts, as they were trained on a limited set of taxa from a single region and have not yet been validated beyond their original geographic or taxonomic scope.

## 2. Results

In addressing this issue, we have developed a novel device named the Edapholog^®^ extractor, which automates the detection and analysis of soil microarthropod diversity. Our previous prototypes based on infrared gate detection were also named the Edapholog probe^29,30^. Traditional methods for collecting soil microarthropods typically involve extracting organisms from soil samples using techniques that exploit their positive geotaxis^36^. In our study, computer vision detected all organisms that passed through the 2 mm mesh in the soil extraction, while the smallest individual detected had a body length of 0.15 mm due to limitations of camera resolution. As a result, our dataset primarily includes microarthropods such as springtails and mites, but also encompasses a variety of other soil-dwelling, wingless arthropods, including e.g., diplurans, symphylans, centipedes, and larvae of flies and beetles. For simplicity, we collectively refer to these organisms as microarthropods^37^. This “dry extraction” approach exploits a vertical moisture gradient to drive microarthropods downward through a sieve, where they would typically be captured in preservative liquids. In contrast, our method preserves the organisms alive. As they move through the sieve into the funnel, they reach a photographic plate, where they remain active during detection. Our system employs video recording combined with a real-time computer vision algorithm. While microarthropods move within the sensing chamber, the algorithm continuously processes the footage, identifying individuals. Once an organism is identified with sufficient confidence, it is blown away from the photographic plate of the sensing chamber into a sample container. The detection and AI-based image analysis of an individual is completed within 0.17 second; if multiple individuals are present, the process runs in parallel. To facilitate rapid image acquisition, we developed a digital microscope paired with a microcomputer optimized for AI modelling (Extended Fig.E1 and Supplementary Methods S1). We employed a computer vision-on-the-edge technology, specifically a MobileNet V2 transfer learning approach embedded within a hyperparameter optimization framework based on the Tree-structured Parzen Estimator (TPE). This integrated method is designed to optimize neural network architectures and hyperparameters systematically, thereby achieving efficient, adaptive, and resource-friendly deep learning models suitable for real-time applications (Extended Fig.E2-E3, Supplementary Methods S2). MobileNet V2 transfer learning has proven effective across various insect classification tasks, applying two-level TPE optimization strategy^33–36^.

Here, we present the results of our AI-driven analyses conducted on arable fields across 46 locations in ten European countries (Fig.1). A total of 319 arable fields were sampled, yielding 34,717 soil microarthropods.

**Figure 1.**
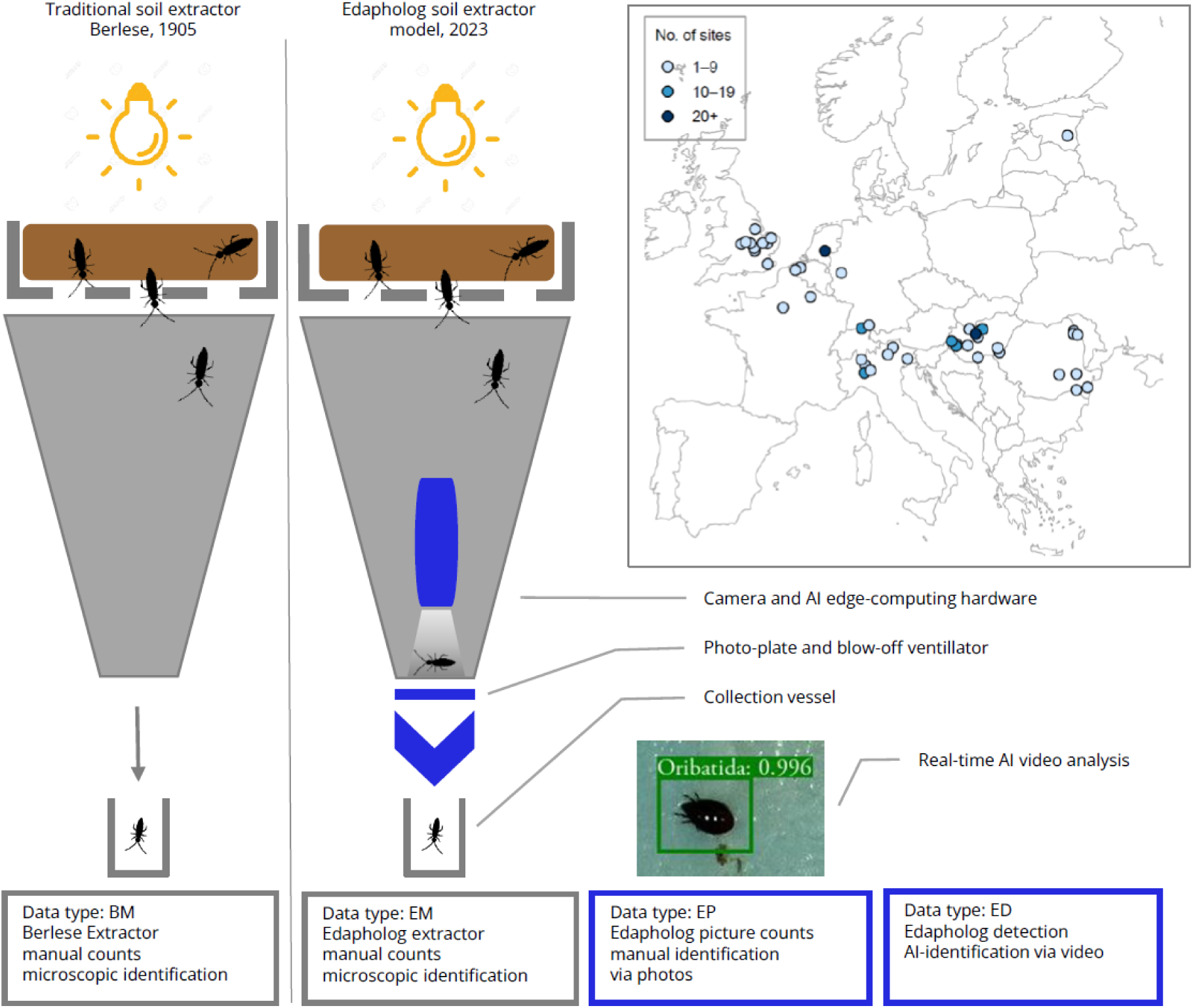
Observational framework for evaluating the accuracy of the new technology. In the classical soil extraction funnel method, microarthropods are extracted from soil and fall into a sample container, where they are killed and preserved (left). Manual microscopic taxonomic identification is then performed, producing taxa-abundance data (BM data). In the new approach (right), the process begins similarly; however, instead of immediate preservation, the living specimens are video-recorded. Motion detection, coupled with a real-time AI-driven algorithm, is used to taxonomically identify them (ED data). After identification, each specimen is blown into a collection vessel, resulting in the same biological sample as the classical method (EM data). Images captured during the automated detection process are stored, allowing for later manual identification (EP data). To validate the new technology, we compared the BM and EM data. The internal biases of the AI system were assessed in two ways. First, we compared the AI-generated data (ED) to the EM data to evaluate detection accuracy; this comparison was performed at the sample level. Then we assessed identification accuracy by comparing the ED and EP data; each organism’s taxonomy was recorded, so this was done at the individual level. By combining the data of all individuals from the same soil sample, we also obtained taxa-abundance data from the EP data. Inset map: 46 sampling locations comprising 319 arable fields across 10 European countries.

We validated the new AI-driven technology by comparing taxa-abundance data obtained using the Edapholog system (ED: AI detection; EM: manual identification of Edapholog-extracted individuals; EP: manual identification from the photographs) with that derived from classical extraction methods (BM; Fig.1). We hypothesized that both approaches would yield similar estimates of mesofaunal diversity from the same community under identical sampling constraints. During detection and transfer (blowing) in the Edapholog, some individuals may escape, potentially introducing bias in the species spectrum. Estimating validity is inherently challenging because a single soil sample can only be processed by one method, requiring comparisons across different but ideally equivalent samples. Given the high spatial heterogeneity and naturally patchy distribution of soil mesofauna, diversity estimates often exhibit high variability^38^. To mitigate this, we applied two approaches. First, in a soil mixing test, multiple soil samples were homogenized and randomly partitioned into smaller, uniform samples. This approach yielded generally high agreement between methods (ED-BM) on a logarithmic scale; however, we observed significant differences in effect sizes for several taxa. (Fig.2A, Supplementary Notes S3.1). Notably, the live-extraction method captured fewer centipedes (*Chilopoda*), millipedes (*Diplopoda*) and pseudoscorpions (*Pseudoscorpiones*), while the AI-driven system performed better for ants (*Formicidae*) and springtails (*Collembola*). The biological samples of the two types of soil extraction (EM-BM) yielded very similar patterns suggesting that taxa-abundance distributions of EM and ED did not differ significantly (Extended Fig.E4A, Supplementary Notes S3.2).

**Figure 2.**
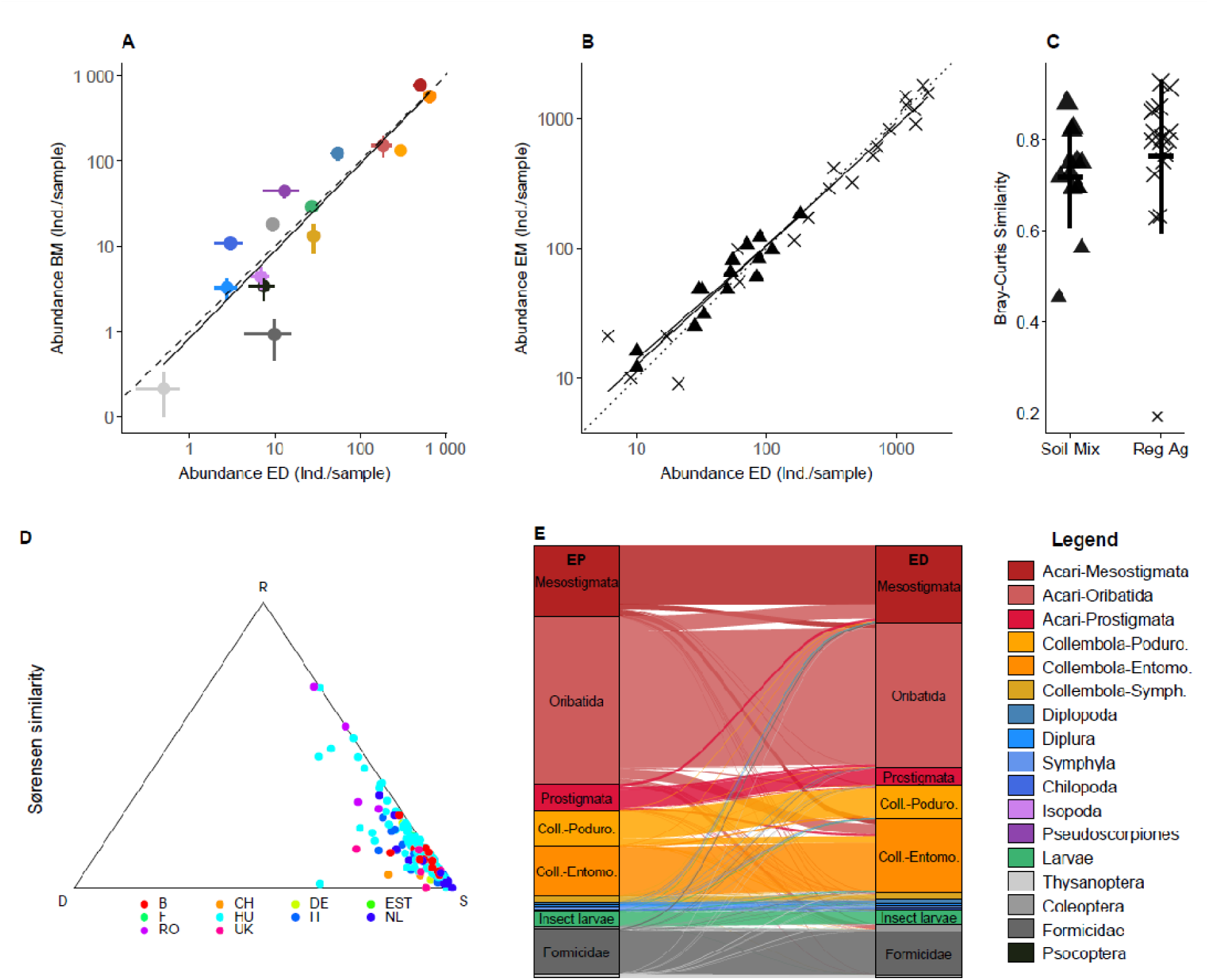
Performance and compositional comparisons of classical and AI-based soil mesofauna extraction and identification. (A) Log-log comparison of mean taxon-level abundances (n = 7 replicates) between Edapholog detection (ED) and classical Berlese extraction (BM) in the Soil-Mix experiment. Points represent taxonomic groups; horizontal and vertical lines show standard errors. The dashed line indicates 1:1 correspondence; the solid line shows the linear fit. (B) Comparison of abundances between ED and EM (manual identification from Edapholog material) across Soil-Mix and Reg-Ag. experiments. Points represent individual samples; shape indicates experimental site. (C) Bray–Curtis similarity between ED and EM compositional data in two experiments. Jittered points show individual soil samples; bars indicate mean ± SD. (D) Sørensen-based SDR triangle plot comparing ED and EP (manual identifications from Edapholog photos) community composition. Each dot represents a plot, 165 plots in ten countries; colours indicate country. (E) Sankey diagram showing identification correspondence between EP (actual class) and ED (predicted class) across 17 microarthropod groups. Widths reflect detection frequency. Internal labels simplify group names; colours correspond to taxonomic identity.

Although dry extraction techniques such as the Berlese and MacFadyen type have been used for over a century^36,39,40^, the current ISO standard lacks detailed technical and metrological specifications^41^. As a result, existing soil extractors, whether they are commercially available or homemade, vary in efficiency. To eliminate design-related bias in estimating absolute population size and species composition, we constructed classical extractors identical in design to the AI-driven system. In the second approach, we collected paired samples from the same plots for direct comparison. This experiment showed greater variability in ED-BM comparison, likely due to spatial heterogeneity and the use of slightly different extractor designs (Extended Fig.E4B, Supplementary Notes S3.3). However, we found high congruence between the biological samples and AI-driven taxa-abundance data generated by the same Edapholog device from the same soil samples (ED-EM). Abundances of the detected microarthropods (ED) and manually counted biological material (EM) correlated well (Fig.2B, Supplementary Notes S4.1) and as did abundance-based community composition (mean Bray-Curtis similarities Soil-Mix = 0.76 and Reg-Ag = 0.71; Fig.2C, Supplementary Notes S4.2). Therefore, the potential bias in life extraction resulting from detection failures and incomplete collection of individuals into conservation vials was likely minimized. Edapholog detected soil microarthropods as they fall through the funnel onto the photographic plate in real time, allowing us to track the temporal progression of soil extractions. By plotting the cumulative number of individuals over time, we observed saturation curves. The asymptotic shape of both individual- and taxon-level curves indicated effective extraction, with no additional detections near the end of the process. The asymptotic plateaus were typically reached well before the end of the extraction period (5 days), except in a few exceptionally wet soil samples where detections persisted longer (Extended Fig.E5).

To assess the accuracy of the AI-driven identification, we compared AI-generated identifications (ED) with manually identified photo records (EP) for 34717 individuals. We achieved an overall accuracy of 86%, a sensitivity (recall) of 75%, and a specificity (true negative rate) of 99% (Fig.2E, Extended Fig.E6, Extended Table E7, Supplementary Notes S6). The proportion of taxa correctly identified by the AI, reflected in an overall recall of 75%, stemmed from two primary sources. First, despite achieving high precision for soil particles (97%), they were overrepresented in misclassified detections (see row and column “Soil” in Extended Fig.E6). This highlights the importance of minimizing soil debris during extraction. Second, identifying mites (Acari)^42^ proved challenging: recall rates for the order Mesostigmata, suborder Oribatida (with Astigmata), and suborder Prostigmata were 78%, 77%, and 66%, respectively (Extended Fig.E8). This challenge could be addressed by (i) accepting class-level identification, as done in the QBS index^43^, or (ii) further subdividing suborders into more refined categories, though our current training dataset size limits this latter approach. Geographic variation in mesofaunal species composition presents a key challenge for AI-based identification. In all countries, average accuracy exceeded 80% and sensitivity (recall) was above 70%, except in Romania, France and Estonia, where sample sizes were too low for reliable estimates (Extended Fig.E9, Supplementary Notes S6).

To demonstrate to community ecologists the impact that taxonomic misidentifications by AI may have on community structure, we compared the community composition of 160 plots (all samples with total abundance > 20 individuals, total number of individuals = 33,724) using AI-driven datasets (ED) and manually identified datasets (EP) from images (Fig.2D). Given the complexity of compositional similarity, we employed the recently introduced SDR-simplex method to visualize the similarity (S), difference (D), and replacement (R) in community composition. The results show that individual sampling points are clustered near the similarity corner of the triangle, indicating that community composition was largely consistent between ED and EP samples. Similarity values exceeded 82% in all countries except Romania (RO) and Estonia (EST). Notably, most of the points falling along the S–R edge of the triangle, with low richness differences (D < 0.07) and minimal replacement (R < 0.15), indicate high agreement, suggesting that very few taxa appeared only in one of the sample types (ED or EP). This finding provides strong evidence that AI does not introduce a bias favouring any particular taxon. When testing for differences in soil community structure among countries based on the three SDR metrics (D, R, and S), we found a significant overall difference (PERMANOVA: F = 3.49, R² = 0.138, p = 0.007), primarily driven by samples from Romania, as confirmed by pairwise PERMANOVA results (Supplementary Notes S5).

To evaluate the utility of the Edapholog device for ecological assessment in arable land, we tested it in a well-documented, 17-year-old field trial comparing conventional and conservation tillage with cover cropping (Reg-Ag. Test, Extended Fig.E5G). Both classical microscopy-based method and the Edapholog system were used to assess mesofaunal responses. As expected, most microarthropod taxa showed increased abundance in conservation tillage plots, with effect sizes ranging from 0.5 to 4 (Fig.3A, CT vs PT). These trends were consistently detected by both classical and AI-based methods (EM vs ED), with effect sizes closely aligned despite the limited replication typical of long-term agricultural field trials (see Supplementary Table S7.1-2 Pairwise contrasts by taxa in Supplementary Notes S7). Micro- and macro-decomposer groups generally responded positively, with all target taxa showing positive effect sizes, except for insect larvae and a small group of non-soil arthropods (e.g., incidental captures such as barklice (*Psocoptera*), which were aggregated into an ‘other’ category outside the primary scope of this study). Oribatid and Mesostigmatid mites, along with major Collembola lineages, exhibited significant increases under conservation tillage. We observed significant shifts in community composition under conservation tillage (Fig.3B, Supplementary Notes S7.2), RDA1 explained 29% of total community variation, indicating a strong separation between tillage types. The total variance explained by the model (Tillage + Time) was 29%, meaning these predictors account for nearly a third of the community structure observed in the ED method data. PERMANOVA confirmed significant differences in community composition between CT and PT (ED: R² = 0.14, p = 0.001; EM: R² = 0.11, p = 0.002).

**Figure 3.**
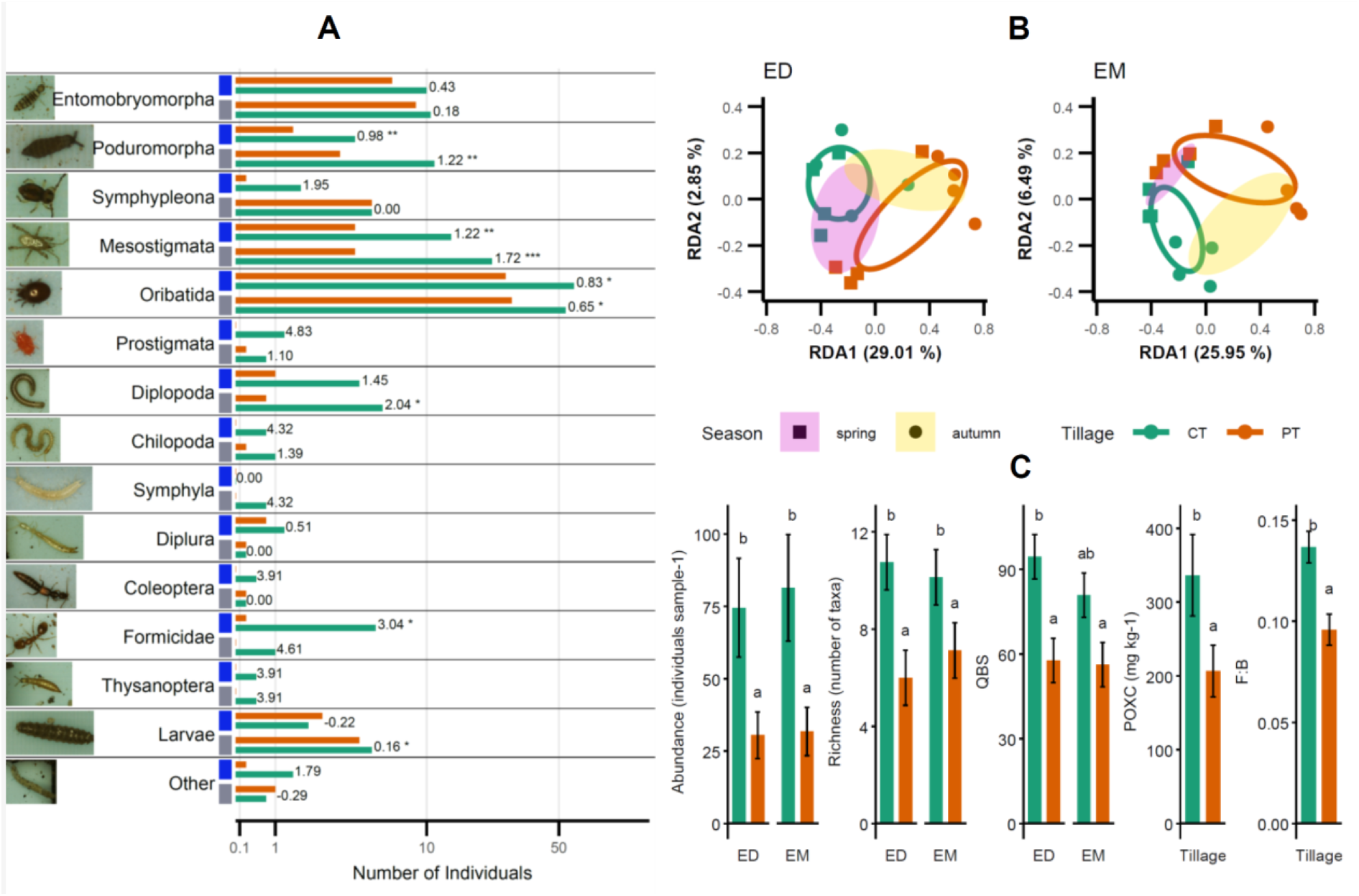
Responses of the microarthropod community to regenerative agricultural practices in a long-term field experiment (Reg-Ag. Test). (A) Mean abundances of major soil microarthropod groups sampled in May under conservation tillage (CT, green) and plough tillage (PT, orange), shown on a logarithmic scale (0.1–50). Blue and grey squares indicate the AI-driven and classic method (blue = ED, grey = EM). Icons represent key taxa detected by the Edapholog system, Effect sizes (log[CT/PT]) and significance markers (p < 0.05, p < 0.01, etc.) indicate differences between tillage types. (B) dbRDA ordination of community composition under CT and PT based on Bray–Curtis distances, shown separately for ED (left) and EM (right). Points represent samples coloured by tillage and shaped by season (spring vs. autumn). Confidence ellipses (95%) and centroid trajectories highlight seasonal and treatment-related shifts in community structure. (C) Estimated marginal means (± SE) of community metrics (abundance, richness, QBS-ar), labile carbon, and the fungal-to-bacterial (F:B) ratio under CT and PT. Bars compare ED (left panels) and EM (right panels), with significance indicated by lowercase letters (different letters = significant difference at p < 0.05).

To integrate community responses into a soil health indicator, we calculated the total abundance, richness and the QBS index, which scores microarthropods based on their morphological adaptation to soil life, under the premise that greater representation of well-adapted forms reflects higher ecological quality^44,43^. To enable automatic QBS calculation, our AI model was trained to recognize the morphological groups used in QBS scoring (Extended Table E10). Total abundance, richness, and the resulting soil biological health index were 39%, 47%, and 150% higher, respectively, under conservation tillage compared to conventional ploughing. Community metrics obtained via manual (EM) and automated (ED) methods showed consistent and significant differences between soil management regimes (Fig.3C). Additionally, to indicate positive effects of further soil biological functions, we measured active soil carbon content and fungal-to-bacterial ratio. Both variables were higher under conservation tillage and correlated strongly with QBS values, as confirmed by GLM models (Supplementary Notes S7.3).

## 3. Discussion

While the positive effects of no-till, mulching, and cover cropping on soil fauna are well established^15,16,45,46^, key challenges remain: how to optimize soil management practices to enhance biodiversity, and how to monitor the impact of regenerative methods reliably and affordably^14^. Here, we addressed the latter through robust, repeatable post-hoc assessments of mesofaunal biodiversity using edge-based computer vision. Further development of the monitoring framework, including sampling frequency and timing, spatial allocation, and data interpretation, will be essential. One may ask whether higher taxonomic resolution is achievable with computer vision–based soil fauna identification. While our training database includes nearly 150 morphologically defined groups, achieving true species–level resolution, particularly within large and morphologically diverse taxa such as Collembola and Acari, remains a challenge, given the taxonomic depth developed over nearly a century. Importantly, soil biodiversity monitoring demands a multidimensional approach: biodiversity ‘sensu stricto’ (i.e. species composition) alone is insufficient, as ecologically distant groups co-exist in functionally integrated soil biomes^47^. This technology offers a novel opportunity to monitor soil ecological processes in situ: notably, the microarthropod individuals remain alive during measurement, enabling downstream experiments on growth, reproduction, and stress tolerance^48^. Moreover, it facilitates a shift from point-based, retrospective assessments to predictive, spatially explicit monitoring systems supported by real-time environmental data streams from platforms such as Google Earth Engine and the Copernicus Sentinel network^49^. Although this technology can be used in any ecosystem type, a key consideration in the system’s design was its applicability to field parcels, as required by carbon farming and precision agriculture. One of the most critical practical needs in these contexts is the ability to integrate soil biodiversity sampling with routine soil analyses, such as nutrient and chemical profiling, particularly with respect to synchronized sampling schedules. Our method is well suited to this integration. In addition to quantifying soil microarthropods, it accurately detects larvae and nymphs, including identifiable forms such as click-beetle larvae and both adult and immature stages of thrips. While integrated pest management already employs forecasting tools, aligning soil biodiversity monitoring with these systems could extend its utility to plant protection, potentially providing strong incentives for farmers to engage in biodiversity assessment as part of carbon farming initiatives.

## 5. Methods

### Computer Vision

We implemented a deep learning pipeline optimized for real-time image classification on our resource-constrained edge device (Supplementary Methods S2). Our approach integrates transfer learning, Bayesian hyperparameter optimization, and quantization, ensuring model efficiency and suitability for deployment^1^. Images were categorized into 24 classes, with 80% used for training and 20% for validation. We applied transfer learning using a pre-trained MobileNet V2, selected for its balance between accuracy, inference speed, and compatibility with TensorFlow 2.x. The MobileNet V2 backbone was used for low-level feature extraction, while upper layers were fine-tuned for the classification task. A two-level hyperparameter optimization (HPO) framework, using the Tree-structured Parzen Estimator (TPE), was applied to jointly optimize architecture and training parameters. In Level 1, various transfer learning configurations were evaluated over a single epoch. Key parameters included the number of frozen layers (20–153), optimizer choice (Adam or SGD)^2^ and learning rates, convolutional kernel size (3×3, 5×5, 7×7), number of channels (4–64), pooling method (average or max), dropout rates (0–0.9), and dense layer presence/size (32–1024). All ranges are detailed in Supplementary Methods S2, with outcomes visualized in Extended Figure E2. This analysis revealed that (1) Layer unfreezing improved performance, particularly with Adam; (2) Average pooling consistently outperformed max pooling; (3) Extra convolutional or dense layers were unnecessary; (4) Third dropout layers had minimal effect without dense layers and could be omitted; (5) Increased flexibility in dropout rates (0–0.99) and unfreezing depth (>10 layers) improved tuning in Level 2. In Level 2, selected architectures were refined through extended training, optimizing dropout rates, learning rates (with cosine annealing), number of frozen/unfrozen layers, and number of training epochs (kept low due to transfer learning). The best configuration achieved 99.31% validation accuracy pre-quantization (Supplementary Methods S2). For edge deployment, we applied post-training quantization using TensorFlow Lite, converting the model to 8-bit integer (int8) format. This reduced model size and inference time with minimal accuracy loss (to 98.69%) while maintaining high class-level recall (Extended Figure E3). Additional quantization methodology is provided in the Supplementary Methods S2.

### Soil-Mix Test

To assess the efficiency of the new extraction method, approximately 64 liters of leaf compost soil were homogenized and divided into sixteen 4-liter samples. Eight samples were processed using classic Berlese extractors and eight with the Edapholog^®^ extractor. Soil extractions lasted five days, after which all specimens were counted and identified under stereo microscope.

### Regenerative Agriculture Long-term Field Test (Reg-Ag. test)

This study used an experimental field originally established in 2003 to study conservation tillage in arable field^3^. The plots are located in western Hungary (46°42’15’’ N, 17°02’50’’ E, 176–206 m a.s.l.) in a hilly region located on a 0%–12% east-facing slope which well represents the region’s most typical soil type, topography, climatic conditions and crop rotation. The regional climate is warm-summer humid continental with a mean annual temperature of 11 °C and mean annual precipitation of 600–700 mm. Soil profiles are eroded on the convex upper part of the slopes, while thick soil sections are typical on the lower concave slopes due to sedimentation. The parent material was loess, and the soil was classified as haplic Luvisols^4^. The average cation exchange capacity was 14.27 meq/100g, and the average base saturation was 76.7% in 2023 in the 0–15 cm soil layer. The 32 ha study area was divided into four pairs of conservation tillage (CT) and ploughed tillage (PT) plots of similar size (∼4 ha) resulting eight plots and four replications. The only difference between the plot pairs was the tillage type, and they received the same treatment in every other aspect (crop rotation and crop type, fertilization, and plant protection). The PT consisted of mouldboard ploughing (25–30 cm depth), harrowing, and seed-bed preparation every year. CT was a non-inversion, ploughless tillage system (using disc and cultivator) with a reduced number of tillage operations where approx. 30 % of the soil surface is left covered by crop residues. In this study area, decreasing degradation of soil organic matter and the closely related increasing biological activity and improved soil structure were found in long-term response to changes in conservation tillage^5^.

Soil samples (400 cm³; 8 cm diameter × 8 cm depth) were collected in spring and autumn 2023 (25 May and 13 November) using a cylindrical soil corer. Each plot was sampled for both traditional extraction and Edapholog technology, with three soil cores per method (1.2 L soil per plot). In both methods, extraction lasted 5 days, after which specimens were preserved in 70% ethanol and identified under a stereomicroscope. While preservation and extraction duration were consistent, the extractors differed slightly in mesh size, container volume, and lighting design.

POXC concentration was examined by the permanganate oxidation method by Weil et al. (2003). A 10 ml of 0.02 M KMnO4 solution was added to a 1 g air-dried soil sample. The soil mixture was then shaken at 125 rpm for 5 minutes. A 200 µl of soil solution and 10 ml of distilled water was added afterward and centrifuged at 3000 rpm for 10 minutes for separating the supernatant and filtrate. A spectrophotometer at 565 wavelength was employed to measure the POXC concentration, defining as the carbon (C) that can be oxidized by KMnO4. To determine the sample KMnO4 concentration, the sample absorbance was compared with a standard curve that ranged from 0.005 to 0.02 mol L–1 KMnO4. Sample POXC was calculated as follows: POXC (mg kg-1) = (0.02 − KMnO4 mol L-1) × 9000 mg C mol-1 × 10.

### Quantification/Measurements of soil bacterial and fungal abundance

Real-time quantitative PCR (qPCR) was performed using the Rotor-Gene^®^ Q platform (Qiagen, Hilden, Germany) to amplify the bacterial 16S rRNA and the fungal ITS regions, and the resulting copy numbers were used as a proxy for microbial abundance. First, total soil DNA was extracted from 300 mg of each composite sample using the DNeasy PowerSoil Pro Kit with the QIAcube system (Qiagen, Hilden, Germany) according to the manufacturer’s instructions, except that mechanical lysis was performed with the TissueLyser LT (Qiagen, Hilden, Germany) at 30/s for 2 min. The quality and concentration of the extracted DNA were then measured using the QIAxpert system (Qiagen, Hilden, Germany) and the Quantus™ Fluorometer (Promega, Madison, USA). For qPCR, we used specific primer sets: Eub338/Eub518 for bacteria and ITS1f/5.8s for fungi, as proposed by Fierer et al. (2005)^6^. Each 20-µl qPCR reaction mixture contained 10 µl Luna Universal qPCR Master Mix (New England Biolabs, Ipswich, USA) containing SYBR Green as a detection dye, 4 µl DNA extract (<16 ng), 0.5 µl of each primer at 10 µM, and 5 µl Milli-Q water. Amplifications were performed in triplicate as follows: 2 min at 95 °C for the initial denaturation, followed by 40 cycles of 15 s at 95 °C (denaturation), 20 s at 53 °C (annealing), and 30 s at 72 °C (extension). No-template controls were also included in each qPCR run. The specificity of the qPCR products was verified by melting curve analysis (60°C to 95°C, in 0.5°C increments). To test for potential PCR inhibition, we selected 10 random samples for a dilution test comparing 10× diluted and undiluted DNA extracts, and no inhibition was detected. For absolute quantification, standard curves were generated using 10-fold serial dilutions of the gBlocks^®^ Gene Fragment, a targeted synthetic oligonucleotide^7^, with concentrations ranging from 10^2^ to 10^8^ copies/μL. The R^2^ values for each standard curve were greater than 0.97 and the amplification efficiency for both genes ranged from 103 to 109 %. The fungal:bacterial ratio, a commonly used descriptor of soil disturbance^8^, was calculated by dividing the ITS gene copy number by the 16S gene copy number.

### International Trial

To evaluate the performance of the Edapholog device and the robustness of our AI classification models across diverse mesofaunal communities, we conducted a large-scale soil biodiversity survey at 46 sites across ten European countries (Belgium, Switzerland, Germany, Estonia, France, Hungary, Italy, Netherlands, Romania, and the UK). Soils were sampled using the traditional soil coring method (five cores per field; total volume 2 L) Approximately 50 Edapholog devices were distributed to farms, research institutes, and universities for use during the trial. Sampling was carried out in spring and autumn of 2023 and 2024. The Edapholog units transmitted measurement data (ED) and backup images (EP) via the internet; in cases of network failure, incomplete data were excluded from the final dataset. Detailed monitoring protocols and site descriptions are provided in online data repository in the folder “Edapholog International Survey”.

### Statistical analyses

AI model accuracy was evaluated using the CVMS framework, with key classification metrics presented in Extended Table S7. For abundance data, we applied a generalized linear mixed-effects model (GLMM) workflow that included model selection and residual diagnostics^9–12^.

Community composition similarity between Edapholog (ED) and manually identified (EP) datasets was assessed by partitioning Sørensen beta-diversity into species replacement (R), richness difference (D), and overall similarity (S = 1 − β_total)^13^. Analyses were restricted to samples with ≥20 individuals in both datasets (N = 160). The resulting D–S–R coordinates were plotted to visualize dominant sources of compositional similarity, color-coded by country.

Differences in soil mesofaunal community composition among countries were tested using PERMANOVA^14,15^, based on Euclidean distances and 999 permutations. Only countries with ≥3 observations were retained. Pairwise comparisons were conducted using pairwise.perm.manova() (RVAideMemoire), restricted to valid group-level comparisons.

The effects of tillage and detection method on five soil health indicators, (1) arthropod abundance, (2) taxonomic richness, (3) QBS-ar index, (4) fungal:bacterial ratio, and (5) labile carbon (lab-C), were tested using GLMMs (glmmTMB v1.1.8^9^, v4.3.2^16^). Fixed effects included tillage regime and detection method (manual (EM) vs. Edapholog(ED)); random intercepts were specified for time and replicate. Response variable distributions were: negative binomial (abundance), Gaussian (richness, QBS-ar, F:B ratio), and Tweedie (lab-C). One extreme outlier was excluded for the F:B model. Estimated marginal means (EMMs) and contrasts were derived using the emmeans package^12^.

Distance-based redundancy analysis (dbRDA) with Bray-Curtis dissimilarities was applied to test for shifts in microarthropod community composition across tillage regimes and sampling times. Constrained ordination was visualized using ggplot2-based methods with permutation-based significance testing^17–20^.

In the extraction efficiency comparison, we analyzed cumulative counts of microarthropods detected in real time by Edapholog (ED). Smoothed curves were fitted using Generalized Additive Models (GAMs; mgcv::gam()) with penalized splines (bs = “ps”, k = 10). Saturation was defined as cumulative detection ≥99.9%, indicating the endpoint of extraction^21^.

All data, R scripts, and output files used in this study, including those for figure generation and statistical analyses, are available at Figshare (10.6084/m9.figshare.29353823)^22^. The repository is organized by figures, extended figures and supplementary text, each in its own folder containing raw input data, R scripts, generated outputs, and a README file with full documentation. The computational environment and package versions used for each analysis are provided via sessionInfo.txt files.

## Acknowledgements

We are deeply grateful to all those who assisted with field sampling, site selection, and conducting measurements using the Edapholog device, including the following individuals: Davide Mariolu (I), Felicitas Bujnoch (D), Lena Klaus (D), Sebastian Funk (D), Lea Theile (D), Niklas Pohl (D), Dénes Besenyői (H), Nicole Galace (B), Eva de Jong (NL), Bas Allema (NL), Thomas Van Loo (B), Edward Van Linden (B), Ruben Mistiaen (B), Mads Falster Husballe (B), Julen Verzeaux (FR), Francois-Xavier Bauer (FT).

## author contributions

The contributions to this article are structured as follows: conceptualization and introduction of the novel construction (B.Z., M.D.); AI model development (B.Z., G.B.); creation of the learning database, including taxonomic identification (V.G., N.F., M.D.); design and carrying out of accuracy assessments (L.S., M.D.); coordination of international monitoring (V.V., M.N.); leading and implementation of field studies in the countries (N.G., T.R.H., V.P., R.K., K.J., A.J., L.S., J.L., E.D., J.B.); statistical analyses (Z.T., D.S., M.D.); critical review and editing of the manuscript (T.R.H., A.V., C.S.);

## competing interest declaration

The authors declare no competing interests.

---

**Extended Figure E1.**
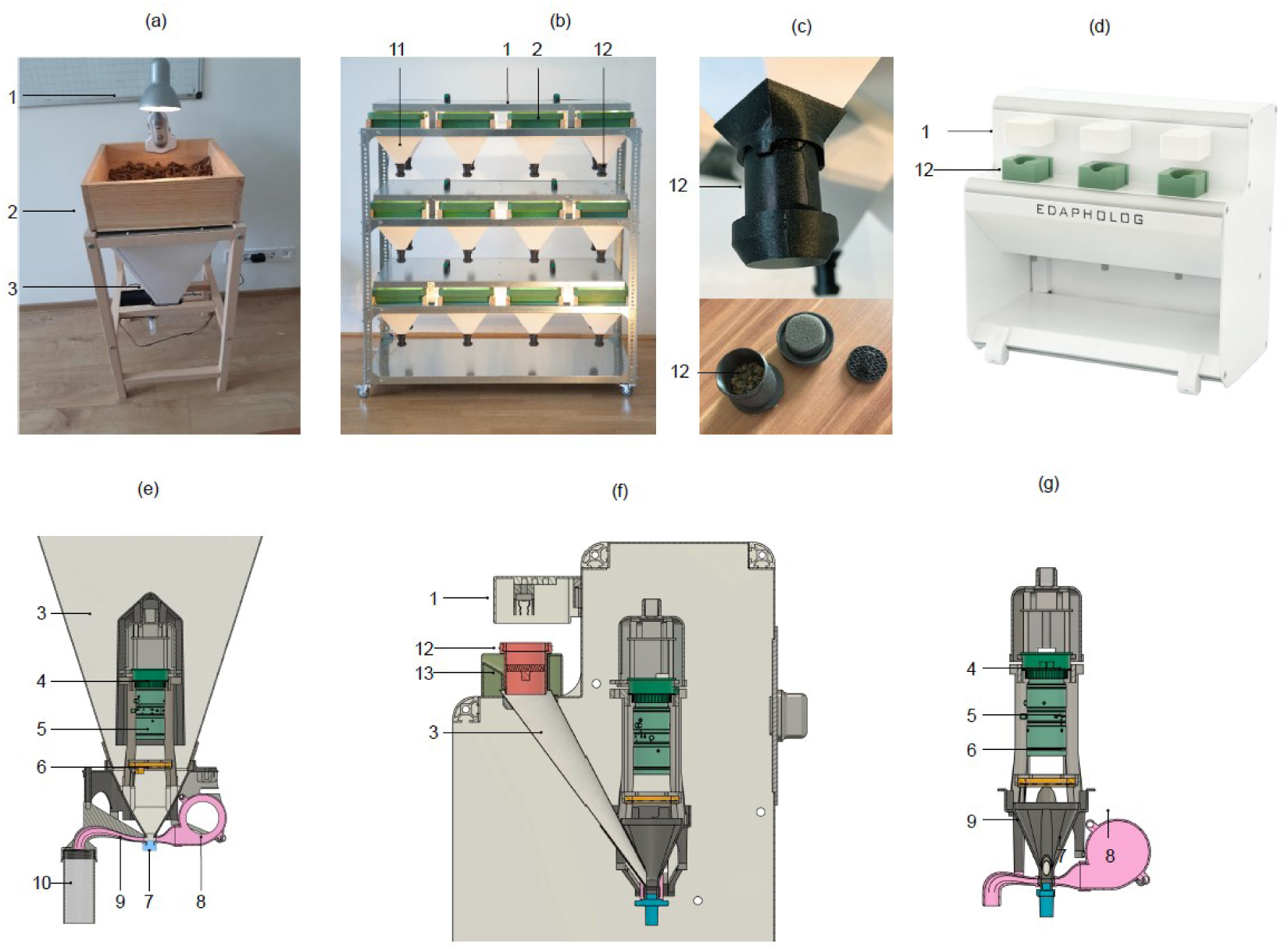
Mechanical design of the Edapholog device. (a): Mechanical construction of the original 2022 model. A 2×2 mm sieve is installed at the base of the soil sample holder (2), targeting mesofauna. A 60° paper funnel (3) directs extracted specimens into the photo chamber while minimizing condensation from soil moisture. A 20 W lamp creates the heat and moisture gradient fundamental to the Berlese–Tullgren method (1). Cross-sectional view (e): A video camera (4), mounted perpendicularly above the photo chamber (7), captures dorsal images of specimens. The camera is enclosed in a smooth-surfaced cover to prevent loss of clinging individuals. The rubber-like material of the photo chamber (7) allows specimen movement, facilitating dorsal imaging. The system uses a Sony IMX477 camera with a Hikrobot MVL-HF3524M-10MP lens (5). The field of view is 8 × 6 mm with a resolution of 2592 × 1944 pixels, yielding 3.08 µm/pixel spatial resolution. A ventilator (8) transfers microarthropods from the photo chamber (7) to the collecting vial (10) via a tunnel (9). Improved version of the device (b–d): The extraction process remains the same, using paper funnels (11) and a heating plate (1), but extracted individuals are collected in a culturing tube (12) (c) instead of being directly imaged. Automatic detection is then performed on organisms extracted from this culturing chamber. This updated design shortens AI-driven detection to under one hour, compared to continuous detection over the full 5-day extraction period. (f–g): Cross-sectional views of the improved design, which retains the core imaging components from the original version. Here, the culturing tube (12) is processed using a small lamp (1) and paper funnel (3). This configuration was used exclusively for validating AI classification accuracy (i.e., ED vs. EP comparisons), as the photographic system remained unchanged. However, faunal composition estimates and all ecological analyses presented in this study were based on the original, fully tested version, since the potential influence of the culturing tube on community composition has not yet been systematically evaluated. The improved design is intended for future experiments involving live individuals; survival tests in the culturing tube will be reported separately.

**Extended Figure E2.**
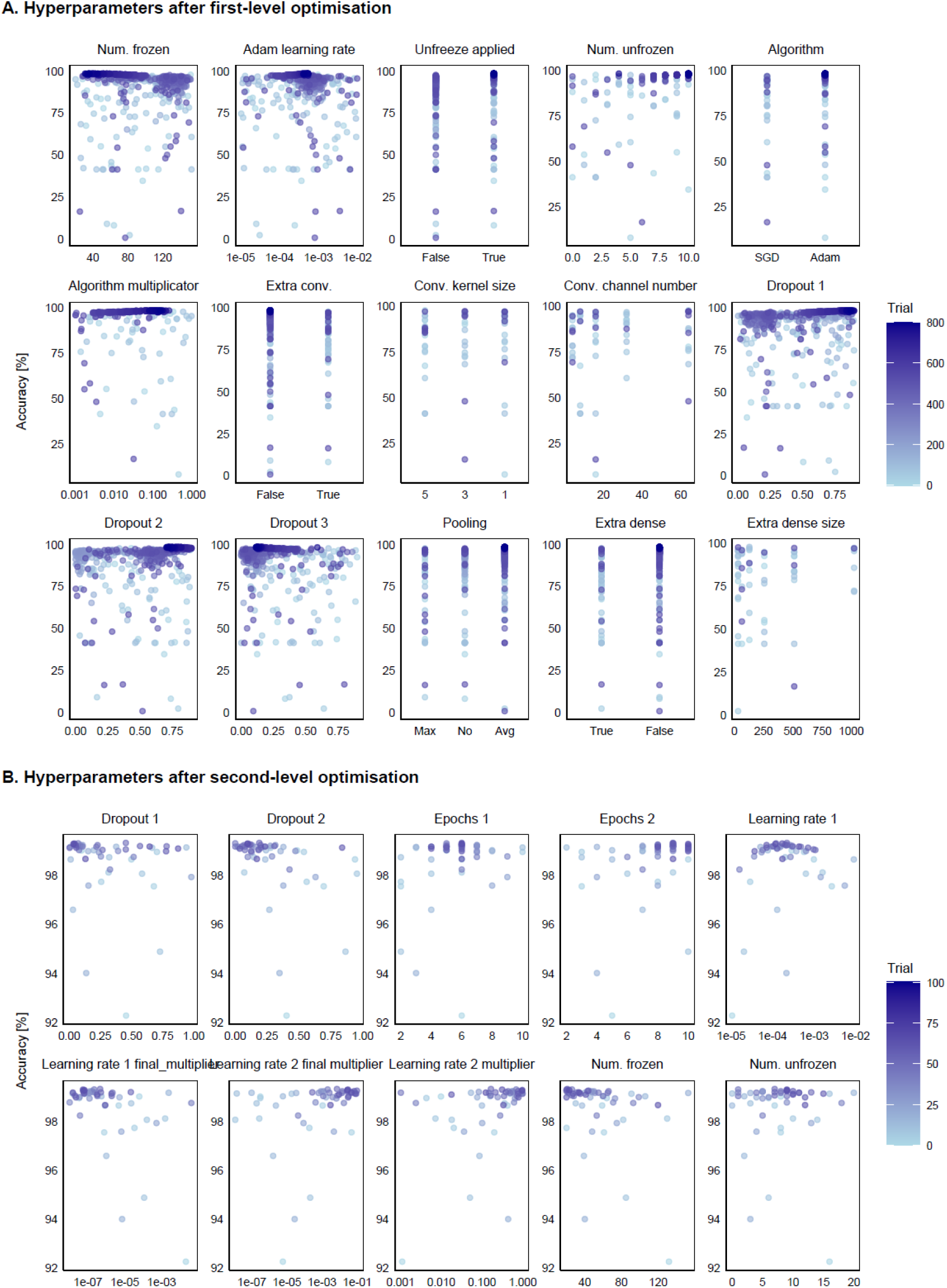
Hyperparameters after the first (A) and second (B) hyperparameter optimization.

**Extended Figure E3.**
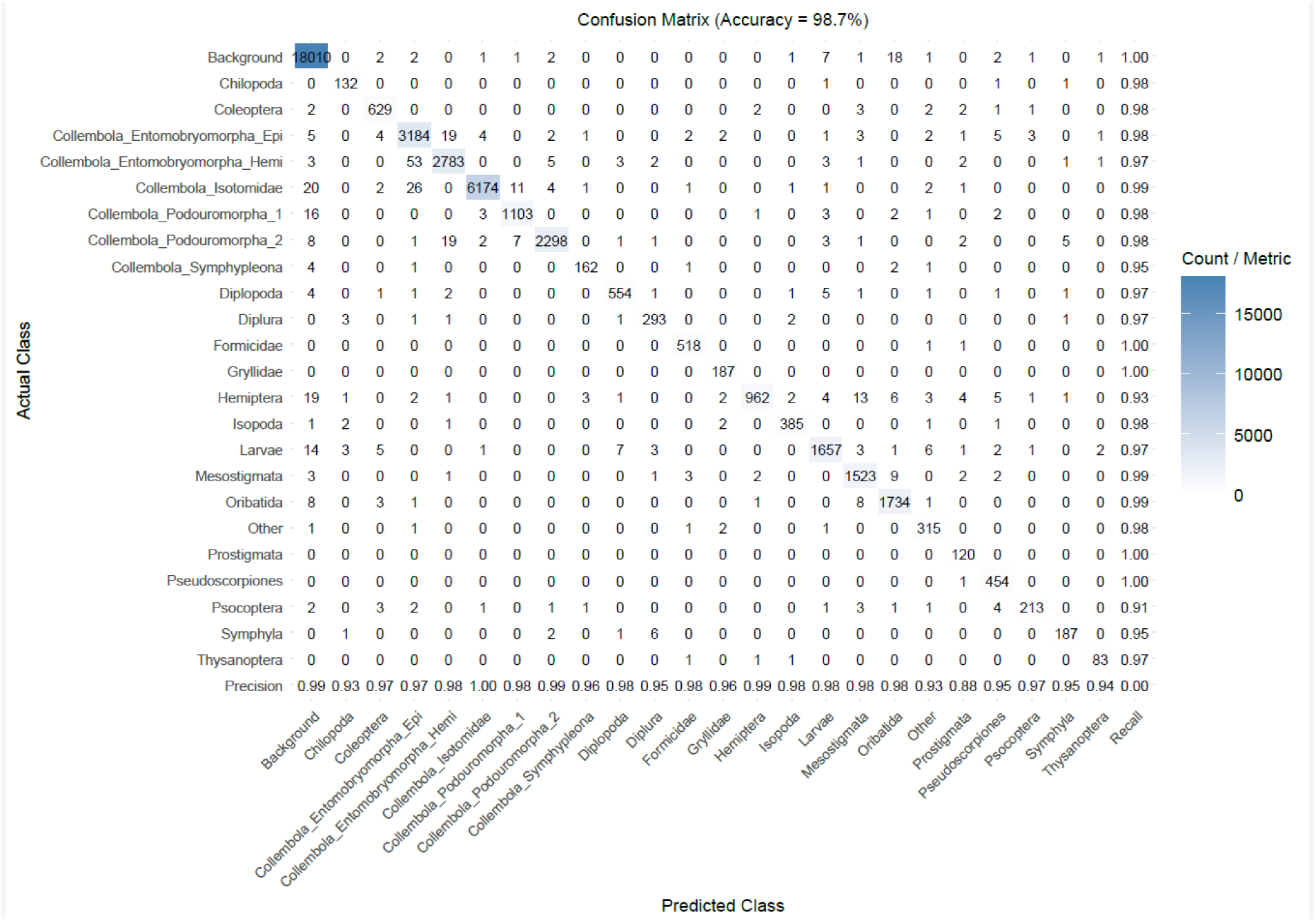
Confusion matrix of the edge model

**Extended Figure E4.**
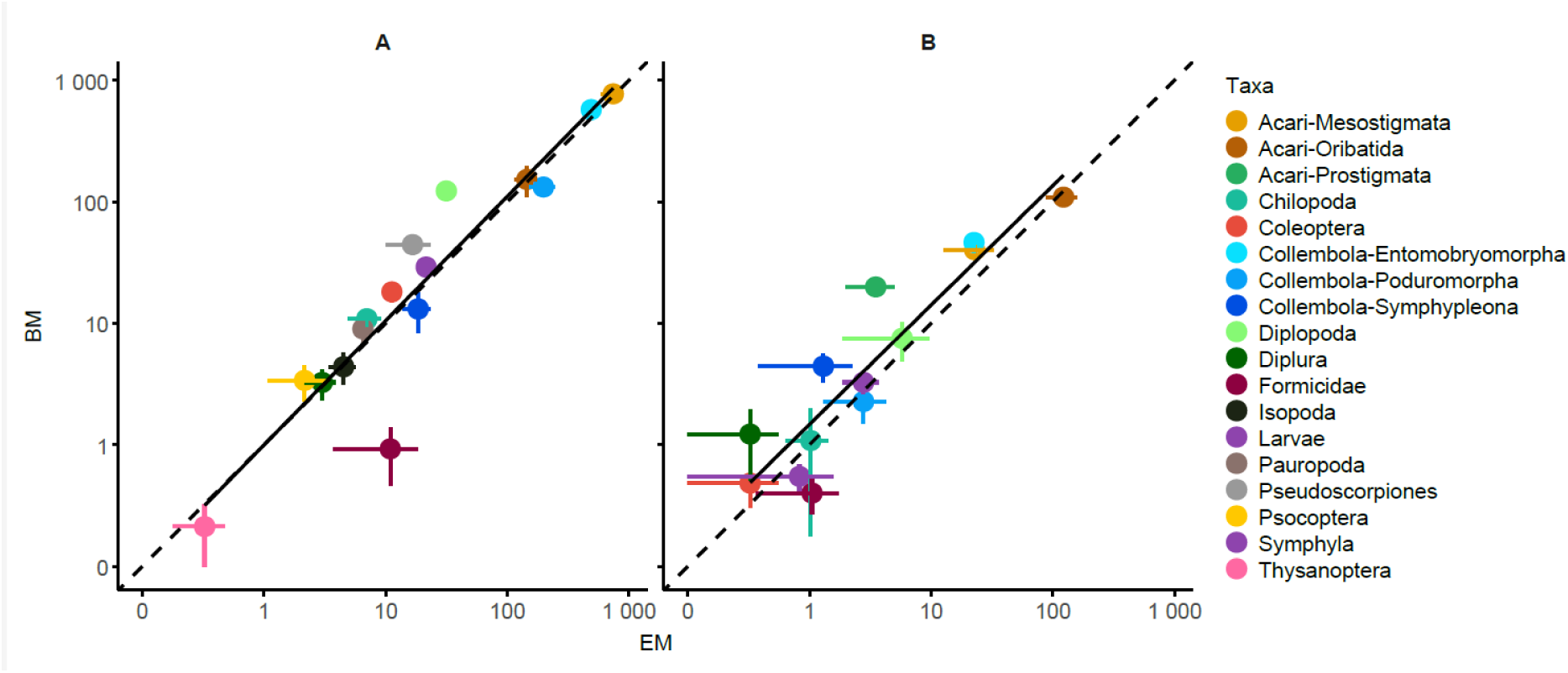
(A) Log-log comparison of mean taxon-level abundances (n = 7 replicates) between manual counting of Edapholog (EM) and classical Berlese extraction (BM) in the Soil-Mix test. Points represent taxonomic groups; horizontal and vertical lines show standard errors. The dashed line indicates 1:1 correspondence; the solid line shows the linear fit. (B) the same comparisons in the Regenerative Agriculture Long-term Field Test (Reg-Ag. test) using replicates from CT and one time point (May). Further statistical test: Supplementary Notes S3.2-3.

**Extended Figure E5:**
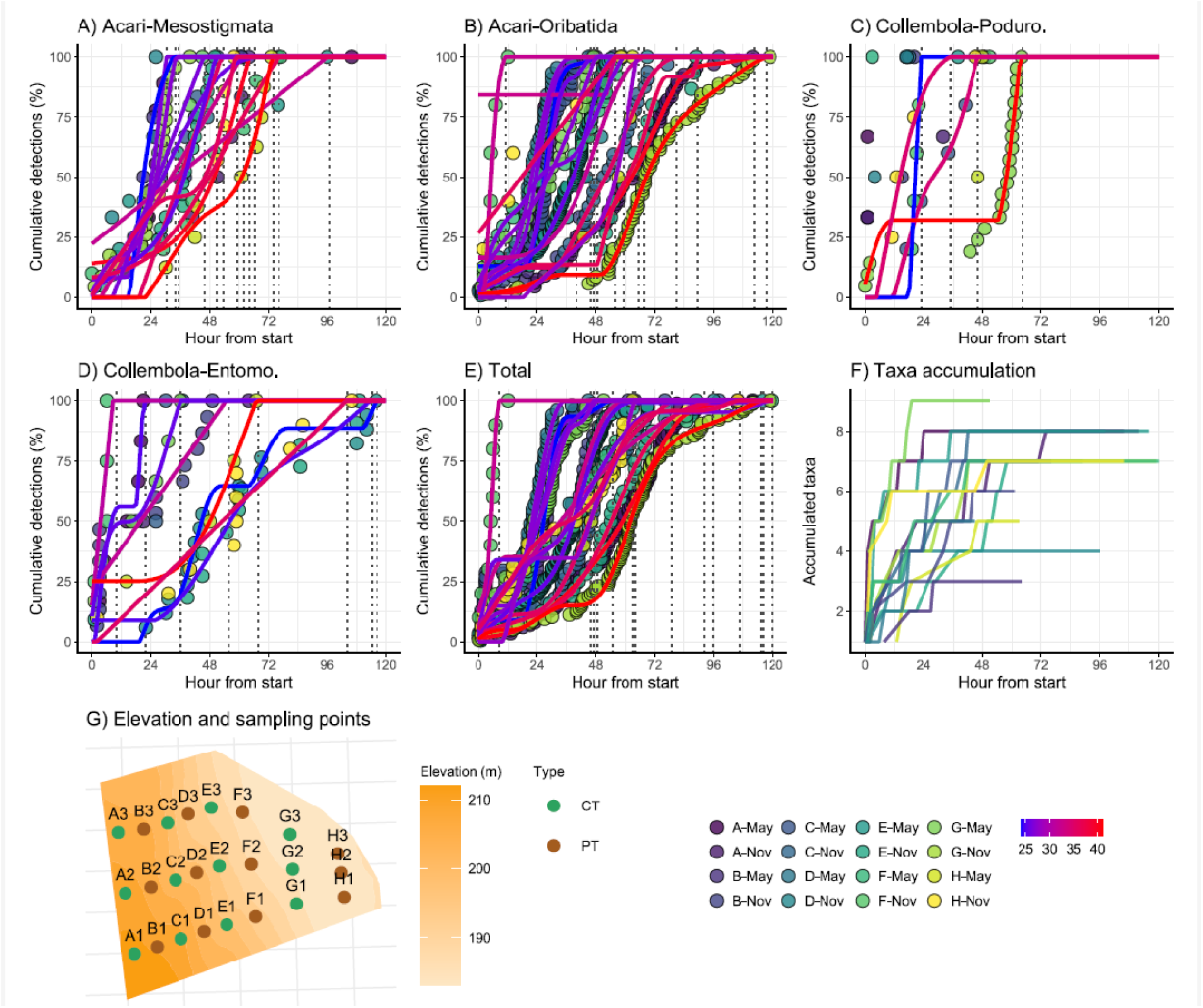
(A–E) Cumulative counts of microarthropods extracted and manually counted (EM) and detected in real time (ED) by the Edapholog device during extraction. Curves represent the most abundant taxonomic groups—Mesostigmata and Oribatida (mites), Poduromorpha and Entomobryomorpha (springtails), and total number of microarthropods. Dot colours indicate the sampling plot at a given time point, while the underlying blue-to-red gradients show initial soil moisture contents (gg-1), smoothed using GAM-based interpolation. (F) Taxon accumulation curves, illustrating the number of newly detected taxa over time. Smoothed curves were fitted using Generalized Additive Models (GAMs). The saturation of both individual and taxon curves indicates that extraction was effective, with no additional detections near the end of the process (F-May and G-May, 119 and 116 hours, initial soil moisture = 0.32 and 0.41gg-1). Plateau was reached when the cumulative detection ≥ 99.9% and remains constant afterward. (G) Plot map (CT: conservation tillage; PT: plough tillage).

**Extended Figure E6.**
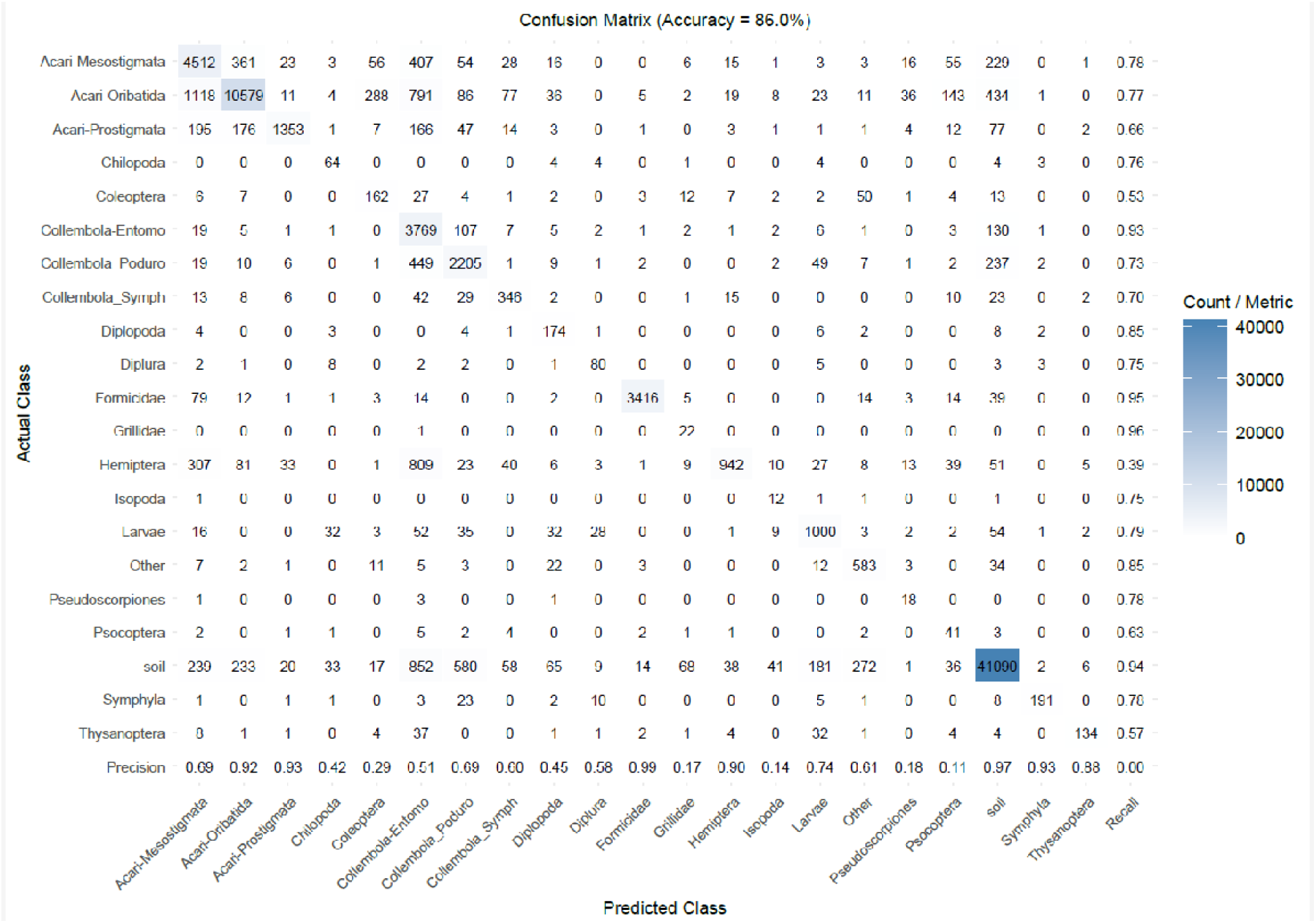
Confusion matrix comparing AI-based predictions (Predicted Class) with manual identifications (Actual Class) of microarthropods. Predicted classes correspond to taxa detected by the AI system (ED), while manual identifications were based on images (EP) of microarthropods captured during the device’s computer vision-based video recordings. Note that in the case of Collembola, the subgroups included in the model developed on the AI learning database were merged into the suborders (Poduromorpha, Entomobryomorpha and Symphypleona). During soil extraction, non-biological particles—such as soil fragments—frequently fall onto the photographic plate where imaging occurs. These must also be classified, and often outnumber true microarthropods, making their exclusion and correct identification a key challenge. Both precision and recall values are presented in the matrix.

**Extended Table E7.** Results of the CVMS analysis, overall metrics.

| CVMS Characteristics | Description | Value |
| --- | --- | --- |
| Overall Accuracy | $\text{Accuracy} = (\text{TP} + \text{TN}) / (\text{TP} + \text{TN} + \text{FP} + \text{FN})$ | 0.86 |
| Balanced Accuracy | $\text{Balanced Accuracy} = (\text{Sensitivity} + \text{Specificity}) / 2$ | 0.87 |
| F1 | $\text{F1} = 2 * \text{Pos Pred Value} * \text{Sensitivity} / (\text{Pos Pred Value} + \text{Sensitivity})$ | 0.62 |
| Sensitivity | Sensitivity (or True Positive Rate, TPR, or recall) = $TP / (TP + FN)$ | 0.75 |
| Specificity | Specificity (or True Negative Rate) = $TN / (TN + FP)$ | 0.99 |
| Pos Pred Value | Pos Pred Value (or Precision) = $TP / (TP + FP)$ | 0.6 |
| Neg Pred Value | Neg Pred Value = $TN / (TN + FN)$ | 0.99 |
| Detection Rate | Detection Rate = $TP / (TP + FN + TN + FP)$ | 0.04 |
| Detection Prevalence | Detection Prevalence = $(TP + FP) / (TP + FN + TN + FP)$ | 0.05 |
| Kappa | For Kappa the counts (TP, TN, FP, FN) are normalized to percentages (summing to 1). Then the following is calculated: $p_{\text{observed}} = TP + TN$; $p_{\text{expected}} = (TN + FP) * (TN + FN) + (FN + TP) * (FP + TP)$;<br>$Kappa = (p_{\text{observed}} - p_{\text{expected}}) / (1 - p_{\text{expected}})$ | 0.61 |

**Extended Figure E8.**
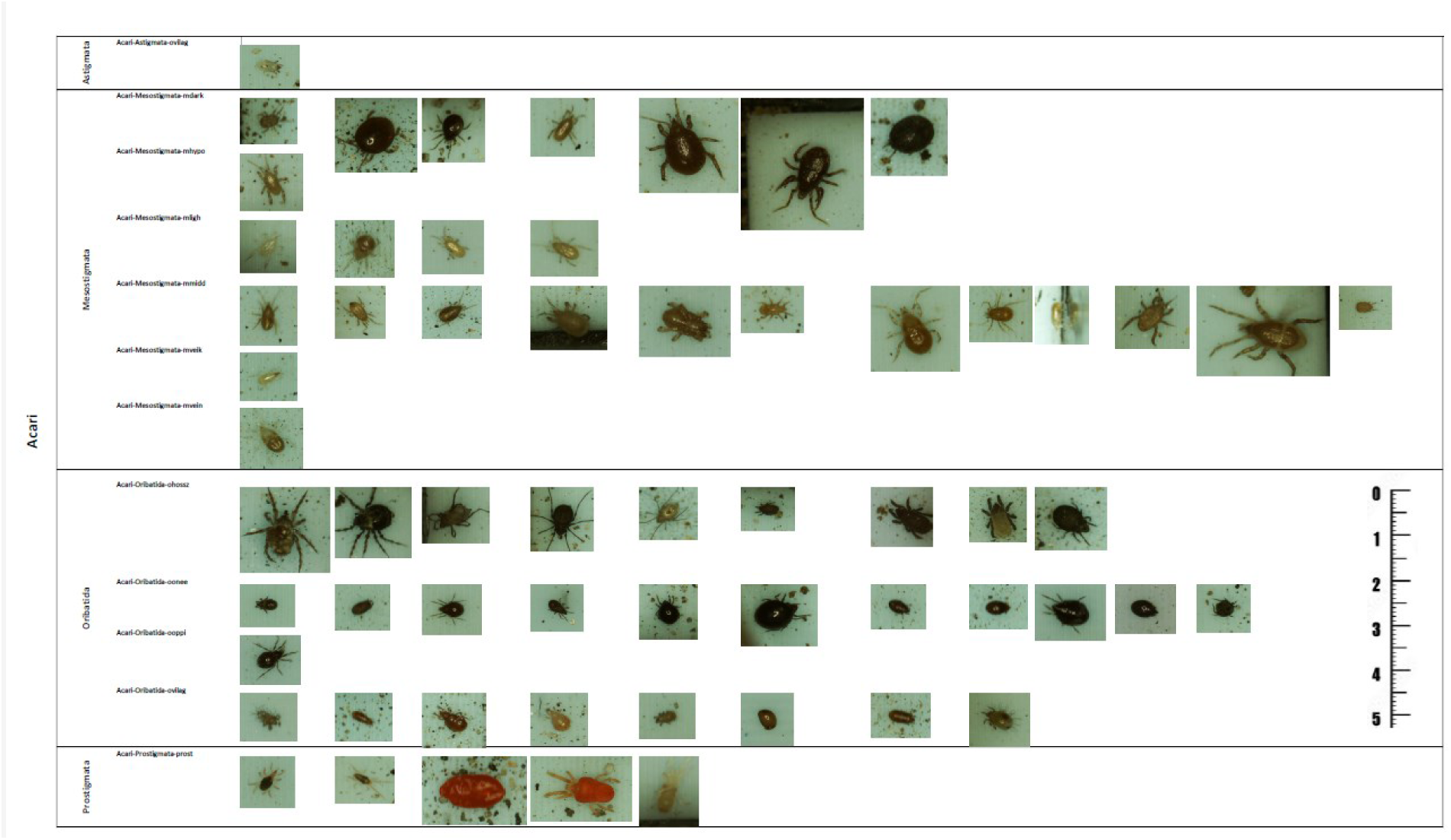
Taxonomy-based grouping strategy for AI model development. Our approach began by identifying all morphologically homogeneous groups that could be distinguished from images, with the support of expert taxonomists. AI models were then trained on these externally uniform groups. Groups were subsequently merged if the model failed to distinguish them reliably, or if too few individual images were available. All images are shown to scale.

**Extended Figure E9.**
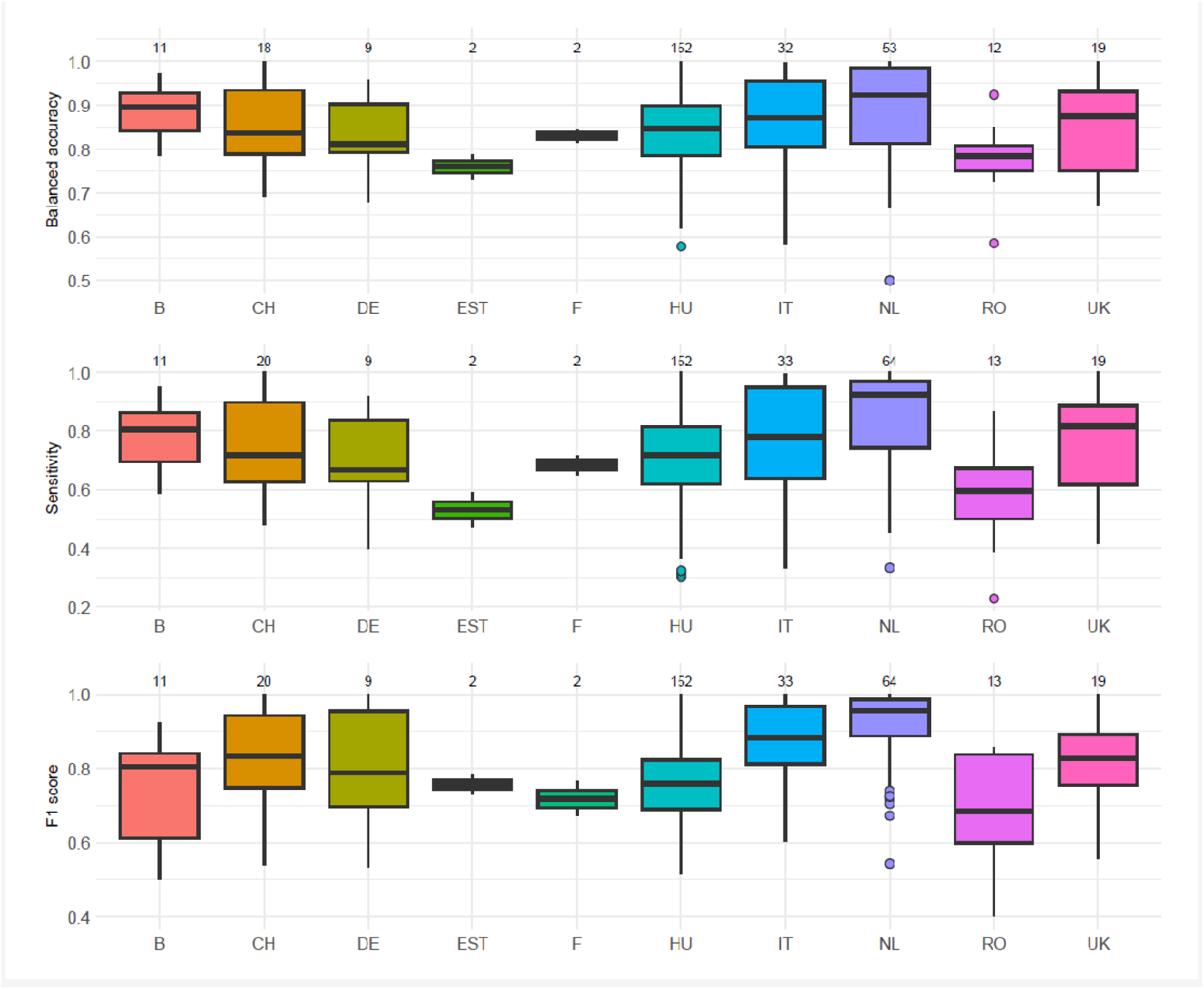
Accuracy of the AI model tested across 10 European countries. Metrics were computed using the CVMS R package. For definitions of the response variables see Extended Table S7. Boxplots: the horizontal line = median, the box = the interquartile range (IQR, Q1–Q3), and whiskers = 1.5×IQR, Outliers = individual points. Sample sizes (number of arable fields) are shown on top.

**Extended Table E10.**
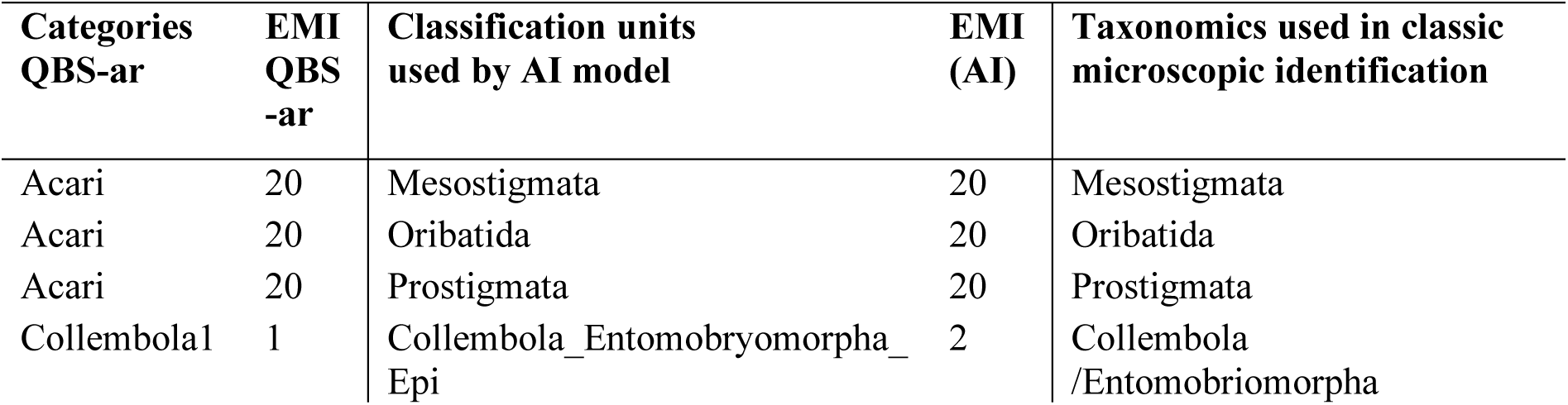
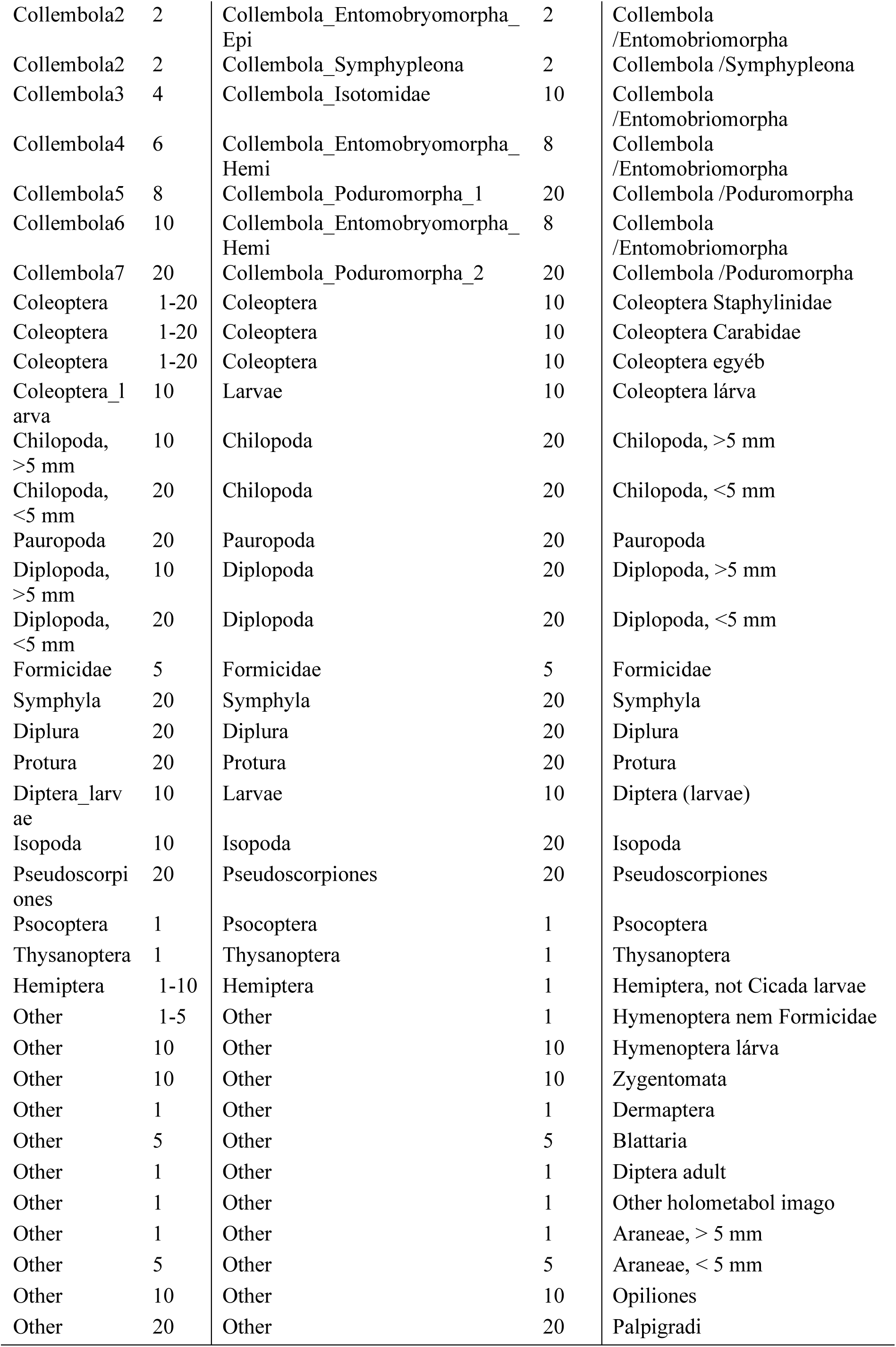

## Notes

https://doi.org/10.6084/m9.figshare.29353823.v1

### Competing Interest Statement

The authors have declared no competing interest.

### Funding

- Co-operative Doctoral Program, HU [C2269690]: KDP-IKT-2023-900-I1-00000957/0000003

