## Supplementary Text for "Edge computer vision produces microarthropod-based high-throughput biodiversity metrics"

### Supplementary Methods S1. Hardware description of the Edapholog® extractor

#### Computer vision

The device continuously records video, during which real-time image analysis is performed on each frame. The image analysis algorithm is presented in Supplementary Figure S1. After the algorithm has identified the animal, the insect is blown from the photo field into the sample container using a ventilator. Image processing consists of the following elements:

a. Motion detection
    a. Background model from history
    b. Subtract current frame
    c. Morphological Transformations
b. Object detection from binary image (cropping)
c. Image classification
d. Detection with statistical operation from multiple images

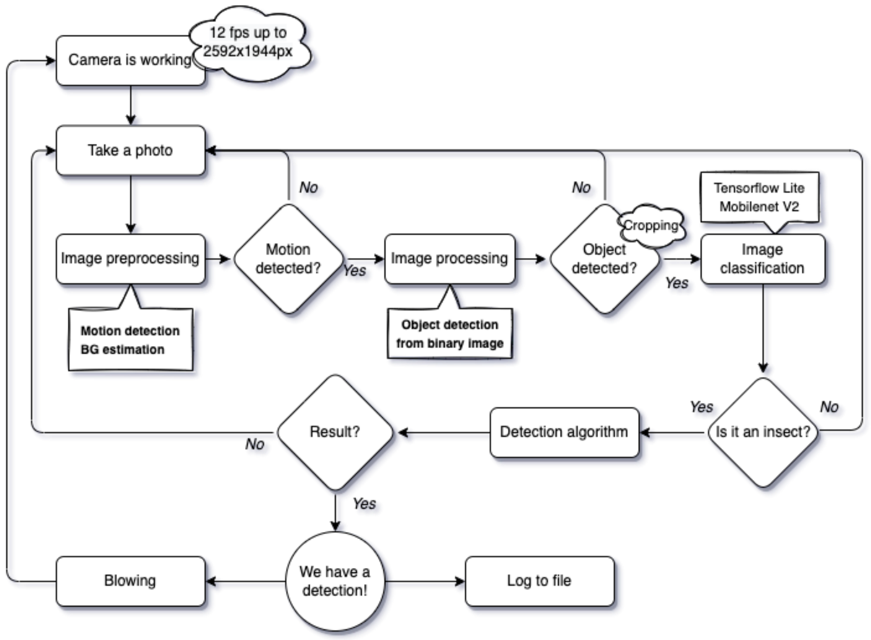

Supplementary Figure S1. Computer vision flow-chart

Supplementary Table S1. Hardware elements of the Edapholog device

| Hardware part | Description |
|---|---|
| Camera | Sony IMX477, 2592 x 1944 (4:3), 20 FPS |
| Lens | Hikrobot MVL-HF3524M-10MP<br>Extension: 20 mm<br>Field of view: 8 x 6 mm<br>Focal length: 35 mm |
| Microcomputer | Coral Dev Board<br>CPU: NXP i.MX 8M SoC (quad Cortex-A53, Cortex-M4F)<br>GPU: Integrated GC7000 Lite Graphics<br>ML accelerator: Google Edge TPU coprocessor: 4 TOPS (int8); 2 TOPS per watt<br>RAM: 1 GB LPDDR4<br>Flash memory: 8 GB eMMC, MicroSD slot<br>Wireless: Wi-Fi 2x2 MIMO (802.11b/g/n/ac 2.4/5GHz) and Bluetooth 4.2<br>USB: Type-C OTG; Type-C power; Type-A 3.0 host; Micro-B serial console<br>LAN: Gigabit Ethernet port |

| | |
|---|---|
| | Audio: 3.5mm audio jack (CTIA compliant); Digital PDM microphone (x2); 2.54mm 4-pin terminal for stereo speakers<br>Video: HDMI 2.0a (full size); 39-pin FFC connector for MIPI-DSI display (4-lane); 24-pin FFC connector for MIPI-CSI2 camera (4-lane)<br>GPIO: 3.3V power rail; 40 - 255 ohms programmable impedance; ~82 mA max current<br>Power: 5V DC (USB Type-C) |
| Control circuit | Custom made board (see. Fig S2)<br>Control circuit runs the microcomputer, led light, blower ventilator. Microcomputer controls actuators via GPIOs.<br>Power: 30 W – 5V(DC) |

The Edapholog's computing unit is a Coral Dev Board, suitable for on-edge AI applications. The camera connects directly to the microcomputer via USB. The system includes a ring light and a blower fan to clear specimens from the photographic plate. Peripheral components are controlled by a custom circuit connected as a HAT via the GPIO pins, enabling software-based control. This board supplies power to the system and includes an RGB LED for feedback. The entire system operates on 5V DC with a maximum power consumption of 30 W.

### Supplementary Methods S2. Description of the Deep Learning Modelling

#### Learning database

The training database was compiled in 2022 using specimens from Hungarian soils and laboratory cultures, independent of the studies presented here. Over 220,000 images were captured with the desktop model of the Edapholog device. The motion detection system recorded multiple frames per specimen in varied positions, enabling more images per individual, especially for rare taxa. However, we used more images from the same specimen mainly for less common groups. Annotation was performed by co-authors using a custom Python tool that streamlined labeling and quality control. This tool also supported validation of experimental datasets by confirming taxa, eliminating low-quality images, and identifying repeated detections of the same individual, which could otherwise bias abundance estimates. All subsequent measurements, including those in this study, can contribute to future model refinement.

A total of 221210 images were collected for the following 24 classes:

- Background also containing soil particles (90250),
- Chilopoda (675),
- Coleoptera (3210),
- Collembola_Entomobryomorpha_Epi (16195),
- Collembola_Entomobryomorpha_Hemi (14285),
- Collembola_ Entomobryomorpha Isotomidae (31220),
- Collembola_Podouromorpha_1 (5655),
- Collembola_Podouromorpha_2 (11740),

- Collembola_Symphypleona (855),
- Diplopoda (2865),
- Diplura (1510),
- Formicidae (2600),
- Gryllidae (935),
- Hemiptera (5150),
- Isopoda (1965),
- Larvae (8530),
- Mesostigmata (7730),
- Oribatida (8780),
- Other (1605),
- Prostigmata (600),
- Pseudoscorpiones (2275),
- Psocoptera (1165),
- Symphyla (985),
- Thysanoptera (430).

As can be seen, the class distribution exhibits a high imbalance ratio of 209.88, calculated as the ratio between the largest and the smallest class. The entropy of the class proportions is 3.16 (on a base-2 logarithmic scale), where lower values indicate more skewed distributions. The maximum entropy for 24 equally distributed classes would be approximately 4.58.

For improved performance in transfer learning, the images were resized to 224×224 pixels. Based on the original image preprocessing method of TensorFlow, each color channel was divided by 127.5 and then 1 was subtracted. An 80 - 20% training-validation split was applied, ensuring that all images of a given specimen appear exclusively in either the training or validation set, but not both.

**Transfer Learning**

Supplementary Table S2-1. Available models for Coral USB Accelerator[1]

| Model name | Input size | Tensorflow version | Latency [ms] | Accuracy [%] | Model size [MB] |
|---|---|---|---|---|---|
| EfficientNet L | 300x300x3 | 1 | 21.3 | 81.2 | 12.8 |
| EfficientNet M | 240x240x3 | 1 | 7.3 | 80.1 | 8.7 |
| EfficientNet S | 224x224x3 | 1 | 5 | 78.9 | 6.8 |
| Inception V1 | 224x224x3 | 1 | 3.4 | 71.9 | 7 |
| Inception V3 | 224x224x3 | 1 | 13.4 | 75.4 | 12 |
| Inception V4 | 299x299x3 | 1 | 84.7 | 80.7 | 42.9 |
| MobileNet V1 | 224x224x3 | 2 | 2.8 | 69.5 | 4.5 |
| MobileNet V2 | 224x224x3 | 2 | 3 | 73.2 | 4.1 |
| ResNet-50 | 224x224x3 | 2 | 42.2 | 73.6 | 25 |

To identify a suitable backbone architecture, we benchmarked various lightweight CNN models compatible with the Coral USB Accelerator (Supplementary Table S1). Extended Figures E2A and B illustrate model trade-offs in latency and size. MobileNet V2 was selected as the optimal choice due to its balance of inference speed, accuracy, small footprint (4.1 MB), and compatibility with TensorFlow 2.x. EfficientNet variants, while competitive, were limited to TensorFlow 1.x and thus excluded for future sustainability. MobileNet V3 was omitted due to incompatibility with the TPU hardware stemming from its use of the Swish[2] activation function[3].

### Hyperparameter Optimization Framework

Supplementary Table S2-2. Search space of first-level hyperparameter optimization

| Number | Hyperparameter | Values |
|---|---|---|
| 1. | Number of frozen layers | 20, 21 … 153 |
| 2. | Adam learning rate | 10-5-0.01 |
| 3. | Unfreeze (second iteration) applied | True, False |
| 4. | Number of unfrozen layers (if 3. is True) | 0, 1 … 10 |
| 5. | Algorithm for unfrozen (if 3. is True) | Adam, SGD |
| 6. | Multiplicator for algorithm (if 3. is True) | 0.001–1 (if 5. is Adam) 0.01–10 (else) |
| 7. | Extra convolution | True, False |
| 8. | Convolution kernel size (if 7. is True) | 3x3, 5x5, 7x7 |
| 9. | Conv. channel number (if 7. is True) | 4, 8, 16, 32, 64 |
| 10. | Dropout 1 | 0–0.9 |
| 11. | Pooling | No, Max, Average |
| 12. | Dropout 2 | 0–0.9 |
| 13. | Extra dense | True, False |
| 14. | Extra dense size (if 13. is True) | 32, 64, 128, 256, 512, 1024 |
| 15. | Dropout 3 | 0–0.9 |

With these configurations, it becomes clear that all previously applied transfer learning methodologies for MobileNet V2[4–7] can be combined. We employed Bayesian optimization via a Tree-structured Parzen Estimator (TPE)[8] to explore the space of architectural and training hyperparameters (Supplementary Table S2-2). The approach enabled efficient evaluation using only two training epochs (1+1), making it suitable for transfer learning scenarios where full retraining is unnecessary.

This first-level optimization revealed:
- Unfreezing layers improved accuracy when using Adam.
- Average pooling outperformed max pooling.
- Extra convolutional/dense layers were unnecessary.
- High dropout rates were beneficial at early layers.
- The third dropout layer could be omitted unless paired with a dense layer.

These insights informed the design of a second-level optimization with a refined parameter space.

### Second-Level Hyperparameter Optimization

Supplementary Table S2-3. Search space of second-level hyperparameter optimization

| Number | Hyperparameter | Values |
|---|---|---|
| 1. | Dropout 1 | 0–0.99 |
| 2. | Dropout 2 | 0–0.99 |
| 3. | Epochs in the first part | 2, 3 … 10 |
| 4. | Epochs in the second part | 2, 3 … 10 |
| 5. | Starting learning rate in the first part | $10^{-5}$–0.01 |
| 6. | Multiplier of the starting learning rate in the second part | 0.001–1 |
| 7. | Multiplier of the finishing learning rate in the first part | $10^{-8}$–0.1 |
| 8. | Multiplier of the finishing learning rate in the second part | $10^{-8}$–0.1 |
| 9. | Number of frozen layers | 20, 21 … 153 |
| 10. | Number of unfrozen layers | 0, 1 … 20 |

The second-level optimization focused on fine-tuning the architecture and training schedule. Dropout values, learning rates (with cosine annealing[8]), layer freezing/unfreezing, and epoch numbers were optimized (Supplementary Table S2-3). Despite fewer iterations, the best configuration reached 99.31% validation accuracy, improving over the 98.06% in the first round. All results exceeded 92%.

The selected configuration featured:
- Medium-length training (6 and 9 epochs for the two phases)
- High dropout in early layers (0.455, 0.190)
- Few frozen (26) and moderate unfrozen (7) layers
- Learning rate schedules starting around $10^{-3}$, decaying via cosine annealing

Full parameter values are in Supplementary Table S2-4.

Supplementary Table S2-4. Best-performing hyperparameter configuration

| Hyperparameter | Values |
|---|---|
| Dropout 1 | 0.455 |
| Dropout 2 | 0.190 |
| Epochs in the first part | 6 |
| Epochs in the second part | 9 |
| Starting learning rate in the first part | $9.788 \times 10^{-4}$ |
| Multiplier of the starting learning rate in the second part | 0.164 |
| Multiplier of the finishing learning rate in the first part | $1.983 \times 10^{-8}$ |
| Multiplier of the finishing learning rate in the second part | 0.0076 |
| Number of frozen layers | 26 |
| Number of unfrozen layers | 7 |

### Quantization

For deployment on low-resource devices, we quantized the trained model to 8-bit integer precision (int8) using TensorFlow Lite. This process reduces model size and computational demand by mapping floating-point weights and activations into the int8 range [-128, 127] via a linear scaling transformation. A representative dataset ensured accurate calibration.

Quantization reduced memory and inference latency with only minor accuracy loss (from

99.31% to 98.69%), while preserving high per-class recall (Figure E5). The model was adapted to batch size 1 for real-time inference on the Coral USB Edge TPU.

**Supplementary Notes S3 (statistics of Figure 2A and Extended Figure E4)**

S3 provides detailed statistical test results to Figure 2A and Extended Figure E4.

**S3.1. Comparison of ED and BM Abundances in the Soil-Mix test, GLM Results (Fig.2A)**

In the Soil-Mix test, we homogenized 64 liters of soil and litter to obtain a standardized mixture containing live soil invertebrates. This mixture was divided into sixteen 4-liter soil samples, of which eight were processed using the classical Berlese method (BM) and eight using the Edapholog device (EM). We tested for significant differences in the mean number of individuals across taxa between the two methods. The 'estimate' column presents effect sizes on a logarithmic scale, they are the log-ratios of abundance between BM and ED. In our model, BM served as the baseline; thus, positive effect sizes (calculated as log(BM) − log(ED)) indicate greater abundance in BM, while negative values indicate greater abundance in EM. To get the 'Ratio of ED to BM'[%], we calculated:

$$\exp(-estimate) = \frac{ED}{BM}*100$$

| Akaike Information Criterion (AIC) | Bayesian Information Criterion (BIC) | Log-Likelihood | Deviance (-2 × log-likelihood) | Residual Degrees of Freedom (df.resid) |
|---|---|---|---|---|
| 1843.2 | 1952.5 | -889.6 | 1779.2 | 193 |

| Response: Abundance | Chisq | Df | Pr(>Chisq) |
|---|---|---|---|
| Method | 0.311 | 1 | 0.577 |
| Taxon | 1,641.809 | 14 | <0.001 |
| Method:Taxon | 74.224 | 14 | <0.001 |

| Taxon | contrast | Ratio_ED to BM | estimate | SE | z.ratio | p.value |
|---|---|---|---|---|---|---|
| Acari-Mesostigmata | BM - ED | 66% | 0.421 | 0.228 | 1.844 | 0.065 |
| Acari-Oribatida | BM - ED | 122% | -0.203 | 0.237 | -0.854 | 0.393 |
| Chilopoda | BM - ED | 27% | 1.308 | 0.466 | 2.805 | 0.005 |
| Coleoptera | BM - ED | 57% | 0.561 | 0.341 | 1.646 | 0.100 |

| | | | | | | |
|---|---|---|---|---|---|---|
| Collembola-Entomobryomorpha | BM - ED | 114% | -0.129 | 0.228 | -0.567 | 0.571 |
| Collembola-Poduromorpha | BM - ED | 225% | -0.811 | 0.236 | -3.437 | 0.001 |
| Collembola-Symphypleona | BM - ED | 237% | -0.864 | 0.320 | -2.699 | 0.007 |
| Diplopoda | BM - ED | 45% | 0.805 | 0.252 | 3.195 | 0.001 |
| Diplura | BM - ED | 83% | 0.184 | 0.525 | 0.351 | 0.726 |
| Formicidae | BM - ED | 518% | -1.645 | 0.640 | -2.573 | 0.010 |
| Isopoda | BM - ED | 161% | -0.477 | 0.435 | -1.096 | 0.273 |
| Larvae | BM - ED | 93% | 0.073 | 0.287 | 0.253 | 0.800 |
| Pseudoscorpiones | BM - ED | 23% | 1.454 | 0.335 | 4.338 | <0.001 |
| Psocoptera | BM - ED | 196% | -0.673 | 0.458 | -1.471 | 0.141 |
| Thysanoptera | BM - ED | 253% | -0.927 | 1.235 | -0.750 | 0.453 |

#### S3.2. Comparison of EM and BM Abundances in the Soil-Mix Test, GLM Results (Extended Figure E4A)

The same procedures as in S3.1 were followed here; however, we compared taxon-abundance data obtained through classical soil extraction (BM) with data identified under a microscope from the biological material collected via Edapholog extractions (EM).

| Akaike Information Criterion (AIC) | Bayesian Information Criterion (BIC) | Log-Likelihood | Deviance (-2 × log-likelihood) | Residual Degrees of Freedom (df.resid) |
|---|---|---|---|---|
| 2040.9 | 2161.5 | - 986.5 | 1972.9 | 222 |

| Response: Abundance | Chisq | Df | Pr(>Chisq) |
|---|---|---|---|
| Method | 7.794 | 1 | 0.005 |
| Taxon | 1,817.219 | 15 | <0.001 |
| Method:Taxon | 48.212 | 15 | <0.001 |

| Taxon | contrast | Ratio_ ED to BM | estimate | SE | z.ratio | p.value |
|---|---|---|---|---|---|---|

| | | | | | | |
|---|---|---|---|---|---|---|
| Mesostigmata | BM - EM | 96% | 0.039 | 0.233 | 0.170 | 0.865 |
| Chilopoda | BM - EM | 62% | 0.474 | 0.370 | 1.282 | 0.200 |
| Coleoptera | BM - EM | 65% | 0.432 | 0.325 | 1.330 | 0.184 |
| Collembola - Entomobryomorpha | BM - EM | 85% | 0.157 | 0.234 | 0.670 | 0.503 |
| Collembola - Poduromorpha | BM - EM | 148% | -0.394 | 0.241 | -1.634 | 0.102 |
| Collembola - Symphypleona | BM - EM | 150% | -0.408 | 0.327 | -1.248 | 0.212 |
| Diplopoda | BM - EM | 26% | 1.341 | 0.263 | 5.106 | <0.001 |
| Diplura | BM - EM | 96% | 0.041 | 0.486 | 0.083 | 0.934 |
| Formicidae | BM - EM | 507% | -1.623 | 0.677 | -2.396 | 0.017 |
| Isopoda | BM - EM | 108% | -0.081 | 0.447 | -0.181 | 0.857 |
| Larvae | BM - EM | 75% | 0.287 | 0.290 | 0.992 | 0.321 |
| Oribatida | BM - EM | 96% | 0.042 | 0.242 | 0.175 | 0.861 |
| Pauropoda | BM - EM | 79% | 0.238 | 0.378 | 0.628 | 0.530 |
| Pseudoscorpiones | BM - EM | 34% | 1.092 | 0.301 | 3.624 | <0.001 |
| Psocoptera | BM - EM | 61% | 0.501 | 0.532 | 0.942 | 0.346 |
| Thysanoptera | BM - EM | 201% | -0.696 | 1.240 | -0.561 | 0.574 |

#### S3.3. Comparison of EM and BM Abundances in Reg-Ag. test, GLM Results (Extended Figure E4B)

Here we used the data from the Regenerative Agriculture Long-term Field Test (Reg-Ag. Test) obtained by classical soil extraction and Edapholog device. We used replicate plots from a given treatment (CT) and time point (May), and compared the data obtained from samples processed using the EM and BM methods. As the experiment included four replicate plots, the sample size for each method was only 4.

| Akaike Information Criterion (AIC) | Bayesian Information Criterion (BIC) | Log-Likelihood | Deviance (-2 × log-likelihood) | Residual Degrees of Freedom (df.resid) |
|---|---|---|---|---|
| 1095.7 | 1189.1 | -519.8 | 1039.7 | 180 |

| Response: Abundance | Chisq | Df | Pr(>Chisq) |
|---|---|---|---|

| | | | |
|---|---|---|---|
| Method | 7.755 | 1 | 0.005 |
| Taxon | 631.706 | 12 | <0.001 |
| Method:Taxon | 18.947 | 12 | 0.090 |

| Taxon | contrast | estimate | SE | df | z.ratio | p.value |
|---|---|---|---|---|---|---|
| Acari-Mesostigmata | BM - EM | 0.615 | 0.331 | Inf | 1.858 | 0.063 |
| Acari-Oribatida | BM - EM | -0.113 | 0.283 | Inf | -0.398 | 0.691 |
| Acari-Prostigmata | BM - EM | 1.674 | 0.486 | Inf | 3.445 | 0.001 |
| Chilopoda | BM - EM | -1.343 | 0.925 | Inf | -1.453 | 0.146 |
| Coleoptera | BM - EM | 0.355 | 1.140 | Inf | 0.312 | 0.755 |
| Collembola /Entomobryomorpha | BM - EM | 0.685 | 0.325 | Inf | 2.105 | 0.035 |
| Collembola /Poduromorpha | BM - EM | -0.150 | 0.592 | Inf | -0.254 | 0.799 |
| Collembola /Symphypleona | BM - EM | 1.213 | 0.738 | Inf | 1.644 | 0.100 |
| Diplopoda | BM - EM | 0.340 | 0.504 | Inf | 0.674 | 0.500 |
| Diplura | BM - EM | 0.816 | 1.115 | Inf | 0.732 | 0.464 |
| Formicidae | BM - EM | -0.700 | 0.869 | Inf | -0.805 | 0.421 |
| Larvae | BM - EM | 0.142 | 0.560 | Inf | 0.254 | 0.800 |
| Symphyla | BM - EM | 0.331 | 1.065 | Inf | 0.311 | 0.756 |

### Supplementary Notes S4 (statistics of Figure 2B, 2C)

#### S4.1. Comparison of ED and EM Abundances in Soil-Mix and Reg-Ag. tests (Figure 2B)

We compared number of individual microarthropods manually counted from biological material (EM) processed by the Edapholog device with those obtained through AI-based detection (ED), using the same soil samples in both tests. In the Soil-Mix dataset, neither the intercept nor the slope of the fitted line deviated significantly from the 1:1 diagonal, indicating strong agreement. Similarly, no significant differences were observed in the Reg-Ag dataset. (see. Figure 2B)

Simultaneous Tests for General Linear Hypotheses for Soil-Mix test

Fit: lm(formula = Abundance_EM ~ Abundance_ED, data = data_Avar)

| | Estimate | Std. Error | t-value | Pr(>\|t\|) |
|---|---|---|---|---|

| | | | | |
|---|---|---|---|---|
| Intercept | -3.92669 | 53.77286 | -0.073 | 0.995 |
| Slope | 0.945 | 0.06267 | -0.878 | 0.55 |

(Adjusted p values reported -- single-step method)

Simultaneous Tests for General Linear Hypotheses for Reg-Ag test

Fit: lm(formula = Abundance_EM ~ Abundance_ED, data = data_Dioskal)

| | Estimate | Std. Error | t-value | Pr(>\|t\|) |
|---|---|---|---|---|
| Intercept | 6.6126 | 7.3712 | 0.897 | 0.519 |
| Slope | 0.9811 | 0.1007 | -0.188 | 0.964 |

(Adjusted p values reported -- single-step method)

#### S4.2. Comparison of ED and EM similarities using Bray-Curtis index in Soil-Mix and Reg-Ag. tests (Figure 2C)

A Shapiro-Wilk test revealed that Bray-Curtis Similarity values deviated from normality in the TAKI_avar group ($p = 0.0002$), but not in the Dioskal group ($p = 0.142$). A Mann-Whitney U test found no significant difference in Bray-Curtis Similarity between the two tests, Soil-Mix test and Reg-AG test ($W = 152$, $p = 0.086$). (see. Figure 2C)

### Supplementary Notes S5. Statistical test of SDR (Figure 2D)

D = richness difference, S = similarity, R = replacement

| Country | D | R | S |
|---|---|---|---|
| B | 0.02 (0.02) | 0.11 (0.03) | 0.86 (0.04) |
| CH | 0.07 (0.06) | 0.05 (0.05) | 0.88 (0.05) |
| DE | 0.02 (0.01) | 0.09 (0.09) | 0.89 (0.09) |
| EST | 0.01 | 0.26 | 0.72 |
| F | 0.02 (0) | 0.14 (0.07) | 0.84 (0.08) |
| HU | 0.03 (0.01) | 0.15 (0.02) | 0.83 (0.03) |
| IT | 0.06 (0.02) | 0.13 (0.04) | 0.82 (0.05) |
| NL | 0.02 (0.01) | 0.08 (0.04) | 0.9 (0.05) |
| RO | 0.04 (0.03) | 0.33 (0.21) | 0.64 (0.2) |
| UK | 0.07 (0.05) | 0.06 (0.05) | 0.87 (0.08) |
| Mean | 0.03 (0.03) | 0.13 (0.07) | 0.83 (0.07) |

Model adonis2(formula = dsr_matrix ~ group_factor, permutations = 999, method = "euclidean"

| | Df | SumOfSqs | R2 | F | Pr(>F) |
|---|---|---|---|---|---|
| | 7 | 0.7437 | 0.13844 | 3.4891 | 0.007 ** |
| Residual | 152 | 4.6281 | 0.86156 | | |
| Total | 159 | 5.3718 | 1.00000 | | |

There is a significant difference in the multivariate DSR space across the 8 countries ($p = 0.007$). The effect size ($R^2 = 0.138$) means that about 13.8% of the variance in DSR space is explained by country-level differences.

Model pairwise.perm.manova(resp, fact, nperm = 999, p.adjust.method = "none", ...)

| | B | CH | DE | HU | IT | NL | RO | UK |
|---|---|---|---|---|---|---|---|---|
| B | x | | | | | | | |
| CH | 0.13 | x | | | | | | |
| DE | 0.683 | 0.729 | x | | | | | |
| HU | 0.368 | 0.165 | 0.339 | x | | | | |
| IT | 0.162 | 0.131 | 0.205 | 0.539 | x | | | |
| NL | 0.328 | 0.43 | 0.953 | **0.03** | **0.044** | x | | |
| RO | **0.014** | 0.122 | 0.103 | **0.006** | **0.032** | **0.004** | x | |
| UK | 0.234 | 0.952 | 0.731 | 0.142 | 0.173 | 0.39 | 0.035 | x |

**Supplementary Notes S6 (Figure 3)**

Supplementary Table S6.1. Pairwise matrix of p-values obtained in GLM analysis of Balanced Accuracy among counties.

| | B | CH | DE | HU | IT | NL | RO | UK |
|---|---|---|---|---|---|---|---|---|
| B | X | | | | | | | |
| CH | 0.96 | X | | | | | | |
| DE | 0.89 | 1.00 | X | | | | | |
| HU | 0.64 | 1.00 | 1.00 | X | | | | |
| IT | 1.00 | 0.99 | 0.95 | 0.48 | X | | | |
| NL | 1.00 | 0.74 | 0.68 | **0.01** | 0.99 | X | | |
| RO | 0.12 | 0.54 | 0.92 | 0.52 | 0.08 | **0.01** | X | |
| UK | 0.94 | 1.00 | 1.00 | 1.00 | 0.98 | 0.65 | 0.58 | X |

Supplementary Table S6.2. Pairwise matrix of p-values obtained in GLM analysis of Sensitivity among counties.

| | B | CH | DE | HU | IT | NL | RO | UK |
|---|---|---|---|---|---|---|---|---|
| B | X | | | | | | | |
| CH | 1.00 | X | | | | | | |
| DE | 0.93 | 0.99 | X | | | | | |
| HU | 0.67 | 0.91 | 1.00 | X | | | | |
| IT | 1.00 | 1.00 | 0.90 | 0.19 | X | | | |
| NL | 0.97 | 0.33 | 0.21 | **<0.01** | 0.59 | X | | |
| RO | **0.05** | 0.08 | 0.72 | 0.21 | **<0.01** | **<0.01** | X | |
| UK | 1.00 | 1.00 | 0.99 | 0.89 | 1.00 | 0.40 | 0.08 | X |

Supplementary Table S6.3. Pairwise matrix of p-values obtained in GLM analysis of F1 Scores among counties.

| | B | CH | DE | HU | IT | NL | RO | UK |
|---|---|---|---|---|---|---|---|---|
| B | X | | | | | | | |
| CH | 0.49 | X | | | | | | |
| DE | 0.95 | 1.00 | X | | | | | |
| HU | 1.00 | 0.39 | 0.99 | X | | | | |
| IT | **0.02** | 0.75 | 0.61 | 0.00 | X | | | |
| NL | **<0.01** | **0.02** | **0.05** | **<0.01** | 0.60 | X | | |
| RO | 0.97 | **0.03** | 0.39 | 0.32 | **<0.01** | **<0.01** | X | |
| UK | 0.66 | 1.00 | 1.00 | 0.65 | 0.58 | 0.01 | 0.06 | X |

### Supplementary Notes 7. Statistics of Figure 3

#### S7.1 Statistics of Figure 3A

We fitted a Generalized Linear Mixed Model (GLMM) using glmmTMB to test the effects of combined treatment (metil, i.e., detection method + tillage) and taxon on microarthropod abundance. (Formula: abundance ~ metil * taxon + (1 | Time)).

Supplementary Table S7.1-1

| Statistic | Value | Description |
|---|---|---|
| AIC | 1604.7 | Akaike Information Criterion – lower values indicate better fit with fewer parameters |
| BIC | 1863.4 | Bayesian Information Criterion – similar to AIC but with a larger penalty for complexity |
| logLik | -740.3 | Log-likelihood – the logarithm of the likelihood function evaluated at the MLEs |
| -2 * log(L) | 1480.7 | Deviance – used for comparing nested models |
| df.resid | 418 | Residual degrees of freedom – number of independent observations minus estimated parameters |

A relatively low AIC and BIC suggest that the model fits the data well with reasonable complexity. The log-likelihood of -740.3 and corresponding deviance (-2 × logLik = 1480.7) can be used to compare this model against nested alternatives: lower deviance means better fit. Taken together, these values suggest the model converged properly, fits the data with moderate complexity, and provides a good basis for inference and prediction.

Supplementary Table S7.1-2 Pairwise contrasts by taxa

| taxon = Acari-Mesostigmata: | | | | | |
|---|---|---|---|---|---|
| contrast | estimate | SE | df | z.ratio | p.value |

| | | | | | | |
|---|---|---|---|---|---|---|
| ED CT - ED | PT | 1.44E+00 | 4.40E-01 | Inf | 3.263 | 0.0061 |
| ED CT - EM | CT | 4.65E-02 | 4.00E-01 | Inf | 0.115 | 0.9995 |
| ED CT - EM | PT | 1.83E+00 | 4.60E-01 | Inf | 4.02 | 0.0003 |
| ED PT - EM | CT | -1.39E+00 | 4.40E-01 | Inf | -3.168 | 0.0084 |
| ED PT - EM | PT | 3.97E-01 | 4.90E-01 | Inf | 0.815 | 0.8475 |
| EM CT - EM | PT | 1.79E+00 | 4.50E-01 | Inf | 3.94 | 0.0005 |

taxon = Acari-Oribatida:

| contrast | | estimate | SE | df | z.ratio | p.value |
|---|---|---|---|---|---|---|
| ED CT - ED | PT | 1.20E+00 | 3.90E-01 | Inf | 3.053 | 0.0121 |
| ED CT - EM | CT | 1.37E-01 | 3.90E-01 | Inf | 0.355 | 0.9847 |
| ED CT - EM | PT | 1.28E+00 | 4.00E-01 | Inf | 3.247 | 0.0064 |
| ED PT - EM | CT | -1.07E+00 | 3.90E-01 | Inf | -2.702 | 0.0347 |
| ED PT - EM | PT | 7.98E-02 | 4.00E-01 | Inf | 0.199 | 0.9972 |
| EM CT - EM | PT | 1.15E+00 | 4.00E-01 | Inf | 2.898 | 0.0197 |

taxon = Acari-Prostigmata:

| contrast | | estimate | SE | df | z.ratio | p.value |
|---|---|---|---|---|---|---|
| ED CT - ED | PT | 2.50E+01 | 1.03E+05 | Inf | 0 | 1 |
| ED CT - EM | CT | 3.13E-01 | 7.10E-01 | Inf | 0.443 | 0.9709 |
| ED CT - EM | PT | 1.95E+00 | 1.14E+00 | Inf | 1.713 | 0.317 |
| ED PT - EM | CT | -2.47E+01 | 1.03E+05 | Inf | 0 | 1 |
| ED PT - EM | PT | -2.31E+01 | 1.03E+05 | Inf | 0 | 1 |
| EM CT - EM | PT | 1.64E+00 | 1.17E+00 | Inf | 1.407 | 0.4948 |

taxon = Chilopoda:

| contrast | | estimate | SE | df | z.ratio | p.value |
|---|---|---|---|---|---|---|
| ED CT - ED | PT | 1.98E+00 | 1.14E+00 | Inf | 1.732 | 0.3071 |
| ED CT - EM | CT | 4.21E-01 | 7.10E-01 | Inf | 0.596 | 0.9334 |
| ED CT - EM | PT | 2.02E+00 | 1.14E+00 | Inf | 1.774 | 0.2861 |
| ED PT - EM | CT | -1.55E+00 | 1.17E+00 | Inf | -1.334 | 0.5412 |
| ED PT - EM | PT | 4.73E-02 | 1.47E+00 | Inf | 0.032 | 1 |
| EM CT - EM | PT | 1.60E+00 | 1.17E+00 | Inf | 1.375 | 0.5152 |

taxon = Coleoptera:

| contrast | | estimate | SE | df | z.ratio | p.value |
|---|---|---|---|---|---|---|
| ED CT - ED | PT | 1.38E+00 | 1.19E+00 | Inf | 1.165 | 0.649 |
| ED CT - EM | CT | 1.43E+00 | 1.19E+00 | Inf | 1.205 | 0.6236 |
| ED CT - EM | PT | 1.43E+00 | 1.19E+00 | Inf | 1.205 | 0.6236 |
| ED PT - EM | CT | 4.73E-02 | 1.47E+00 | Inf | 0.032 | 1 |
| ED PT - EM | PT | 4.73E-02 | 1.47E+00 | Inf | 0.032 | 1 |
| EM CT - EM | PT | -7.00E-06 | 1.47E+00 | Inf | 0 | 1 |

taxon = Collembola-Entomobryomorpha:

| contrast | | estimate | SE | df | z.ratio | p.value |
|---|---|---|---|---|---|---|
| ED CT - ED | PT | 6.40E-01 | 4.20E-01 | Inf | 1.513 | 0.4297 |

| | | | | | | |
|---|---|---|---|---|---|---|
| ED CT - EM | CT | 1.98E-01 | 4.10E-01 | Inf | 0.478 | 0.9639 |
| ED CT - EM | PT | 7.81E-01 | 4.20E-01 | Inf | 1.839 | 0.2548 |
| ED PT - EM | CT | -4.42E-01 | 4.30E-01 | Inf | -1.04 | 0.7258 |
| ED PT - EM | PT | 1.41E-01 | 4.40E-01 | Inf | 0.324 | 0.9883 |
| EM CT - EM | PT | 5.83E-01 | 4.30E-01 | Inf | 1.368 | 0.5192 |

taxon = Collembola-Poduromorpha:

| contrast | | estimate | SE | df | z.ratio | p.value |
|---|---|---|---|---|---|---|
| ED CT - ED | PT | 1.87E+00 | 5.50E-01 | Inf | 3.414 | 0.0036 |
| ED CT - EM | CT | -1.04E+00 | 4.20E-01 | Inf | -2.462 | 0.0659 |
| ED CT - EM | PT | 3.65E-01 | 4.50E-01 | Inf | 0.809 | 0.8502 |
| ED PT - EM | CT | -2.91E+00 | 5.30E-01 | Inf | -5.449 | <.0001 |
| ED PT - EM | PT | -1.51E+00 | 5.60E-01 | Inf | -2.707 | 0.0343 |
| EM CT - EM | PT | 1.40E+00 | 4.30E-01 | Inf | 3.239 | 0.0066 |

taxon = Collembola-Symphypleona:

| contrast | | estimate | SE | df | z.ratio | p.value |
|---|---|---|---|---|---|---|
| ED CT - ED | PT | 1.06E+00 | 7.00E-01 | Inf | 1.518 | 0.4264 |
| ED CT - EM | CT | -3.35E-01 | 5.30E-01 | Inf | -0.628 | 0.9231 |
| ED CT - EM | PT | -8.45E-01 | 5.20E-01 | Inf | -1.628 | 0.3628 |
| ED PT - EM | CT | -1.40E+00 | 6.70E-01 | Inf | -2.073 | 0.1621 |
| ED PT - EM | PT | -1.91E+00 | 6.60E-01 | Inf | -2.88 | 0.0208 |
| EM CT - EM | PT | -5.09E-01 | 4.80E-01 | Inf | -1.052 | 0.7186 |

taxon = Diplopoda:

| contrast | | estimate | SE | df | z.ratio | p.value |
|---|---|---|---|---|---|---|
| ED CT - ED | PT | 1.19E+00 | 6.10E-01 | Inf | 1.966 | 0.2011 |
| ED CT - EM | CT | -2.23E-01 | 4.80E-01 | Inf | -0.462 | 0.9674 |
| ED CT - EM | PT | 1.36E+00 | 6.30E-01 | Inf | 2.149 | 0.1377 |
| ED PT - EM | CT | -1.41E+00 | 6.00E-01 | Inf | -2.37 | 0.0828 |
| ED PT - EM | PT | 1.70E-01 | 7.20E-01 | Inf | 0.235 | 0.9954 |
| EM CT - EM | PT | 1.58E+00 | 6.20E-01 | Inf | 2.538 | 0.0543 |

taxon = Diplura:

| contrast | | estimate | SE | df | z.ratio | p.value |
|---|---|---|---|---|---|---|
| ED CT - ED | PT | 4.91E-01 | 8.30E-01 | Inf | 0.591 | 0.9348 |
| ED CT - EM | CT | 1.56E+00 | 1.17E+00 | Inf | 1.343 | 0.5353 |
| ED CT - EM | PT | 1.56E+00 | 1.17E+00 | Inf | 1.343 | 0.5353 |
| ED PT - EM | CT | 1.07E+00 | 1.22E+00 | Inf | 0.879 | 0.8156 |
| ED PT - EM | PT | 1.07E+00 | 1.22E+00 | Inf | 0.879 | 0.8156 |
| EM CT - EM | PT | -7.00E-06 | 1.47E+00 | Inf | 0 | 1 |

taxon = Formicidae:

| contrast | | estimate | SE | df | z.ratio | p.value |
|---|---|---|---|---|---|---|
| ED CT - ED | PT | 2.91E+00 | 1.10E+00 | Inf | 2.659 | 0.0392 |
| ED CT - EM | CT | 9.27E-01 | 5.90E-01 | Inf | 1.582 | 0.3892 |

| | | | | | | |
|---|---|---|---|---|---|---|
| ED CT - EM | PT | 2.55E+01 | 8.05E+04 | Inf | 0 | 1 |
| ED PT - EM | CT | -1.99E+00 | 1.14E+00 | Inf | -1.743 | 0.3014 |
| ED PT - EM | PT | 2.26E+01 | 8.05E+04 | Inf | 0 | 1 |
| EM CT - EM | PT | 2.46E+01 | 8.05E+04 | Inf | 0 | 1 |

taxon = Larvae:

| contrast | | estimate | SE | df | z.ratio | p.value |
|---|---|---|---|---|---|---|
| ED CT - ED | PT | 9.50E-01 | 5.20E-01 | Inf | 1.84 | 0.2545 |
| ED CT - EM | CT | -5.92E-01 | 4.50E-01 | Inf | -1.307 | 0.5583 |
| ED CT - EM | PT | 6.31E-01 | 4.90E-01 | Inf | 1.284 | 0.573 |
| ED PT - EM | CT | -1.54E+00 | 5.00E-01 | Inf | -3.094 | 0.0106 |
| ED PT - EM | PT | -3.18E-01 | 5.30E-01 | Inf | -0.601 | 0.9319 |
| EM CT - EM | PT | 1.22E+00 | 4.70E-01 | Inf | 2.588 | 0.0476 |

taxon = Other:

| contrast | | estimate | SE | df | z.ratio | p.value |
|---|---|---|---|---|---|---|
| ED CT - ED | PT | 1.74E+00 | 1.15E+00 | Inf | 1.511 | 0.4307 |
| ED CT - EM | CT | 6.65E-01 | 8.10E-01 | Inf | 0.821 | 0.8447 |
| ED CT - EM | PT | 3.87E-01 | 7.60E-01 | Inf | 0.512 | 0.9562 |
| ED PT - EM | CT | -1.07E+00 | 1.22E+00 | Inf | -0.879 | 0.8156 |
| ED PT - EM | PT | -1.35E+00 | 1.19E+00 | Inf | -1.139 | 0.6654 |
| EM CT - EM | PT | -2.77E-01 | 8.60E-01 | Inf | -0.322 | 0.9884 |

taxon = Symphyla:

| contrast | | estimate | SE | df | z.ratio | p.value |
|---|---|---|---|---|---|---|
| ED CT - ED | PT | 2.06E+01 | 2.07E+04 | Inf | 0.001 | 1 |
| ED CT - EM | CT | -3.04E-01 | 1.00E+00 | Inf | -0.305 | 0.9901 |
| ED CT - EM | PT | 1.82E+01 | 6.24E+03 | Inf | 0.003 | 1 |
| ED PT - EM | CT | -2.09E+01 | 2.07E+04 | Inf | -0.001 | 1 |
| ED PT - EM | PT | -2.40E+00 | 2.16E+04 | Inf | 0 | 1 |
| EM CT - EM | PT | 1.85E+01 | 6.24E+03 | Inf | 0.003 | 1 |

taxon = Thysanoptera:

| contrast | | estimate | SE | df | z.ratio | p.value |
|---|---|---|---|---|---|---|
| ED CT - ED | PT | 2.04E+01 | 1.95E+04 | Inf | 0.001 | 1 |
| ED CT - EM | CT | 4.00E-06 | 1.08E+00 | Inf | 0 | 1 |
| ED CT - EM | PT | 1.80E+01 | 5.81E+03 | Inf | 0.003 | 1 |
| ED PT - EM | CT | -2.04E+01 | 1.95E+04 | Inf | -0.001 | 1 |
| ED PT - EM | PT | -2.42E+00 | 2.04E+04 | Inf | 0 | 1 |
| EM CT - EM | PT | 1.80E+01 | 5.81E+03 | Inf | 0.003 | 1 |

Results are given on the log (not the response) scale.
P value adjustment: Tukey method for comparing a family of 4 estimates

**S7.2 Statistics of Figure 3B**

Supplementary Table S7.2-1 Results of the distance-based redundancy analysis (dbRDA) in Figure 3B.

| Method | RDA1 Percent[1] | RDA2 Percent[1] | Constrained_var[2] | Total_var[3] | Explained_percent[4] | Predictors |
|---|---|---|---|---|---|---|
| ED | 29.01 | 2.85 | 0.853 | 2.679 | 31.9 | Tillage + Time |
| EM | 25.95 | 6.49 | 1.028 | 3.17 | 32.4 | Tillage + Time |

[1]The percentage of the total variance in the species (community) data explained by the first (RDA1) and the second canonical axis (RDA2).

[2]The amount of variance in community composition that is explained by the predictors (e.g., Tillage + Time).

[3]The total variance in the community data, including both explained (constrained) and unexplained (residual).

[4]The proportion of total variance explained by your predictor variables, a core measure of model fit.

Supplementary Table S7.2-1 PERMANOVA summary Table on dbRDA output data

| Method | Comparison | $R^2$ | p-value |
|---|---|---|---|
| ED | Tillage + Time | 0.325 | 0.005 |
| ED | Tillage (May) | 0.193 | 0.224 |
| ED | Tillage (Nov) | 0.378 | 0.028 |
| EM | Tillage + Time | 0.329 | 0.001 |
| EM | Tillage (May) | 0.194 | 0.151 |
| EM | Tillage (Nov) | 0.382 | 0.029 |

**S7.3 Statistics of Figure 3C**

Supplementary Table S7.3-1 Summary table of all model results used in Figure 3C analysis.

| Response | Family | Formula | AIC | Estimate | SE | p-value |
|---|---|---|---|---|---|---|
| F_B | gaussian (identity) | F_B ~ Tillage | -168.4 | -0.04 | 0.01 | <0.000 |
| lab_C | tweedie (log) | lab_C ~ Tillage | 580.3 | -0.49 | 0.18 | 0.006 |
| QBSar | gaussian (identity) | QBSar ~ Tillage * Method | 294.4 | -36.75 | 9.75 | <0.000 |
| Richness | gaussian (identity) | Richness ~ Tillage * Method | 155.3 | -4.75 | 1.07 | <0.000 |
| Abundance | nbinom1 (log) | Abundance ~ Tillage * Method | 279.4 | -0.89 | 0.23 | <0.000 |

Effect: TillagePT

Supplementary Table S7.3-2 Pairwise comparisons in Figure 3C analysis

| Response variable | Method | CT-PT Estimate | SE | p-value |
|---|---|---|---|---|
| Abundance | ED | 0.894 | 0.231 | 0.0001 |
| | EM | 0.94 | 0.227 | <0.0001 |
| Richness | ED | 4.75 | 1.07 | 0.0002 |
| | EM | 3 | 1.07 | 0.0096 |
| QBSar | ED | 36.8 | 9.75 | 0.0008 |
| | EM | 24.6 | 9.75 | 0.018 |

Supplementary Table S7.3-3 Effect of Tillage between ED and EM in Figure 3C analysis

| Response | Interaction Term | Estimate | p-value |
|---|---|---|---|
| QBSar | TillagePT:MethodEM | +12.1 | 0.379 |
| Richness | TillagePT:MethodEM | +1.75 | 0.249 |
| Abundance | TillagePT:MethodEM | –0.045 | 0.889 |

We applied generalized linear mixed models (GLMMs) to assess the effects of tillage regime and detection method on five response variables. All models included "Time" as a random intercept, and fixed effects were assessed using Wald chi-square tests. Significance was defined at $p < 0.05$.

Fungal:Bacterial Ratio (F_B)

A Gaussian model showed a significant negative effect of plough tillage (PT) compared to conservation tillage (CT) (Estimate = –0.041 ± 0.011, p = 0.00019, AIC = –168.4), indicating reduced fungal dominance under PT.

Labile Carbon (lab_C)

A Tweedie model revealed a significant decrease in labile carbon under PT (Estimate = –0.488 ± 0.178, p = 0.006, AIC = 580.3). This suggests reduced carbon availability in conventionally tilled soils.

QBS-ar Index (QBSar)

A Gaussian model with interaction between Tillage and Method showed a strong main effect of tillage (Estimate = –36.75 ± 9.75, p = 0.00016, AIC = 294.4). Pairwise comparisons confirmed lower QBS-ar scores under PT for both methods (CT–PT = 36.8, p = 0.0008 for ED; 24.6, p = 0.018 for EM).

Richness

Richness was significantly reduced by PT (Estimate = –4.75 ± 1.07, p = 9.5e–06, AIC = 155.3). Interaction effects were not significant. Pairwise contrasts showed reductions in both ED and EM methods (CT–PT = 4.75, p = 0.0002 for ED; 3.00, p = 0.0096 for EM).

Total Abundance

A negative binomial model indicated a strong negative effect of PT on arthropod abundance (Estimate = –0.894 ± 0.231, p = 0.00011, AIC = 279.4). Both detection methods supported this pattern (CT–PT = 0.894, p = 0.0001 for ED; 0.940, p < 0.0001 for EM), indicating consistent decline in abundance under conventional tillage.
